# Early development of male germ cell clones shapes their reproductive success

**DOI:** 10.1101/2025.09.02.672339

**Authors:** Tatsuro Ikeda, Maurice Langhinrichs, Tamar Nizharadze, Chieko Koike, Yuzuru Kato, Katsushi Yamaguchi, Shuji Shigenobu, Kana Yoshido, Shinnosuke Suzuki, Toshinori Nakagawa, Ayumi Maruyama, Seiya Mizuno, Satoru Takahashi, Nils B. Becker, Hans-Reimer Rodewald, Thomas Höfer, Shosei Yoshida

**Affiliations:** Division of Germ Cell Biology, National Institute for Basic Biology; Okazaki, 444-8787, Japan; Division of Theoretical Systems Biology, German Cancer Research Center; Heidelberg, 69120, Germany; Japan Society for the Promotion of Science; Tokyo, 102-0083, Japan; Graduate Institute for Advanced Studies, SOKENDAI; Okazaki, 444-8787, Japan; Department of Gene Function and Phenomics, Mammalian Development Laboratory, National Institute of Genetics; Mishima, 411-8582, Japan; Department of Genome Biology, Graduate School of Medicine, Osaka University; Osaka, 565-0871, Japan; School of Biological and Environmental Sciences, Kwansei Gakuin University; Sanda, 669-1330, Japan; Trans-Omics Facility, National Institute for Basic Biology; Okazaki, 444-8787, Japan; Laboratory of Theoretical Biology, Graduate School of Biostudies, Kyoto University; Kyoto, 606-8315, Japan; Laboratory Animal Resource Center in Transborder Medical Research Center, University of Tsukuba; Tsukuba, 305-8575, Japan; Division of Cellular Immunology, German Cancer Research Center; Heidelberg, 69120, Germany

## Abstract

Germ cells secure the continuity and evolutionary potential of species. Yet how reproductive success is allocated among the founder cells of the mammalian male germline, the primordial germ cells (PGCs), remains unknown. Here, we quantitatively traced individual PGCs, non-invasively barcoded in mouse embryos, across their establishment, lifelong maintenance within the testis, spermatogenesis, and transmission to the next generation. We found that highly skewed clonal contributions arise from a very early stochastic bottleneck during migration to the developing testes and remain stable throughout adult reproductive life. Reproductive success during adulthood was proportional to the embryonically established PGC clone size. Mathematical modeling shows that the patchy compartmentalization of PGC clones within the extended thin seminiferous tubules maintain clonal diversity and safeguard against the expansion of clones that gain a selective advantage. These findings uncover fundamental principles governing the development, stability, and evolutionary transmission of the mammalian germline.

## Introduction

The germline, unlike the somatic lineages supporting bodily functions, is passed on to the next generation and thus ultimately governs species continuity and evolution. As in somatic tissues, we expect variation in cellular fitness in germ cells through mutations, epimutations or phenotypic plasticity. Indeed, it has been shown that positive selection can occur in the human male germline, including in spermatogonial stem cells (SSCs)^1^. Elucidating the quantitative rules governing SSC development, spermatogenesis, and, crucially, transmission to the next generation is pivotal for understanding how cellular behaviors within and across generations shape evolution.

In mouse embryos, the germline is specified as a cluster of primordial germ cells (PGCs) in the posterior extraembryonic region around embryonic day (E)6.5. These cells then migrate through the hindgut to colonize the bilateral gonads by E10.5–11.5^2,3^. In the embryonic testis, PGCs are specified to a male fate, differentiate into gonocytes (or prospermatogonia), and are partitioned among 10–11 distinct testis cords that later develop into seminiferous tubules^4^. Gonocytes proliferate mitotically until around E14.5, followed by cell cycle arrest until birth^5,6^. After birth, gonocytes resume proliferation, and a subset establishes the SSC pool that sustain life-long sperm production^7^. These processes have been well characterized at the cell-population level, but the dynamics of individual PGCs and their lineages remain poorly understood. Previous studies employing a limited number of clonal markers have found evidence of variability between PGC clones with respect to proliferation, localization, and cell death^8–11^. To overcome this scale limitation, we conducted a comprehensive PGC clone lineage-tracing analysis, using *in situ Polylox* DNA barcoding^12^, from the development and maintenance of the male germline to its transmission to the next generation.

## Results

### Tracing the PGC-clonal repertoire through development and adulthood

To barcode embryonic PGCs using *Polylox*, we developed a *Prdm14-Mer-iCre-Mer* (*MiCM*) transgenic mouse line, which expresses MiCM, a strictly tamoxifen-dependent Cre recombinase^13,14^ under the control of the *Prdm14* regulatory region (Figures S1A and S1B). *Prdm14* is essential for PGC specification and germline establishment, showing specific expression in PGCs from E6.5 to E14.5^15^. We crossed male and female mice harboring *Prdm14-MiCM* and *Polylox*, respectively, to obtain double-transgenic embryos (Figure 1A and Table S1). By injecting 4-hydroxytamoxifen (4-OHT) into pregnant females, barcodes were generated exclusively in the embryos’ PGCs but not in gonadal somatic cells in a strict 4-OHT dependent manner (Figures S1C–S1E and S2A), without affecting the distribution and number of PGCs (Figure S1F and S1G, and Table S2).

**Figure 1.**
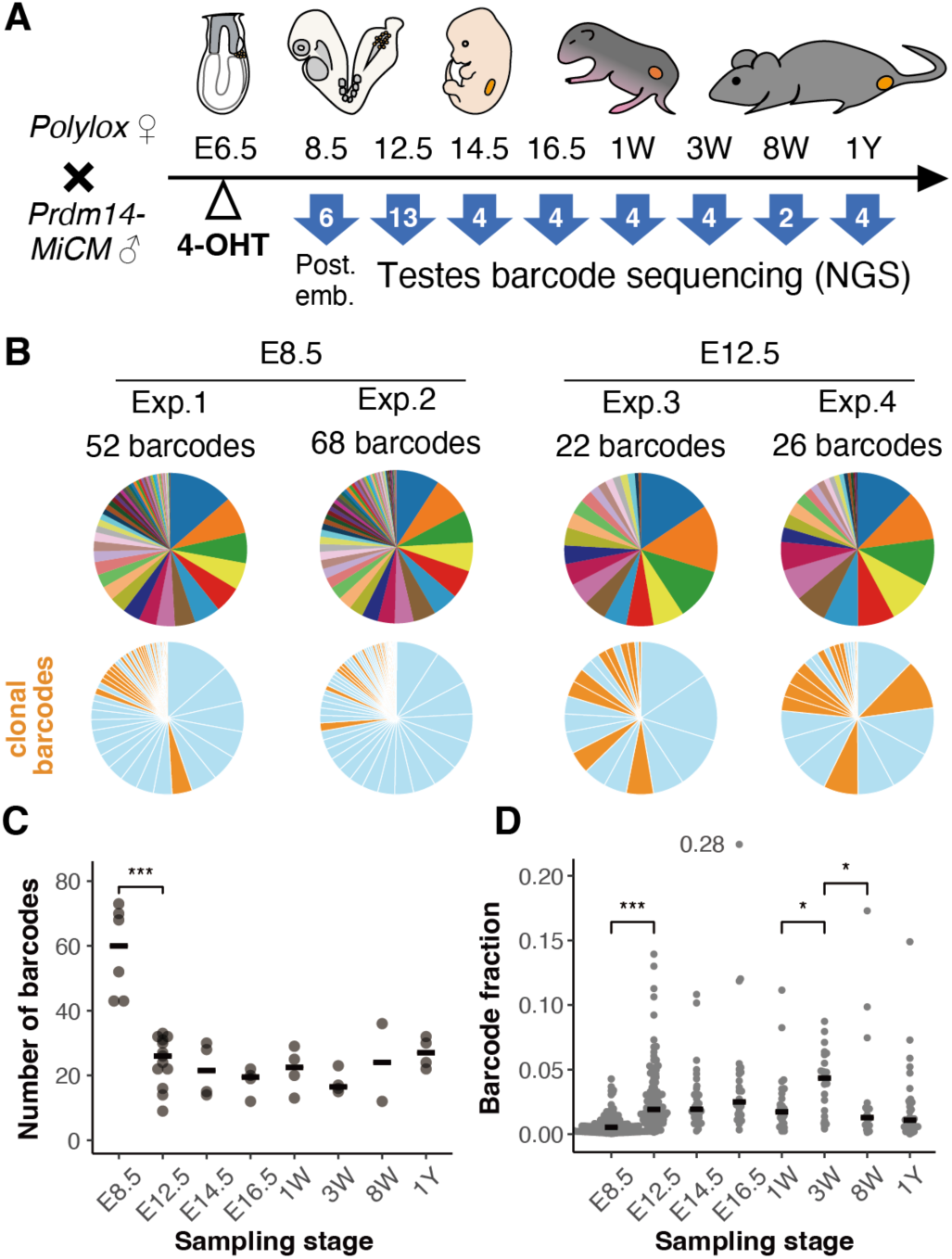
Barcode analysis of E6.5-labeled PGC clones. (**A**) Experimental overview. *Prdm14-MiCM*;*Polylox* male embryos were barcoded at E6.5, whose barcodes in germline were analyzed at the indicated time points from individuals of indicated numbers shown in arrows. Week (W); year (Y). The barcode reads obtained from the testes of 1Y-old mice were from experiments detailed in Figure 2. (**B**) Representative distribution of barcode reads found in two embryos each at E8.5 and E12.5, displayed in pie charts. Unrecombined barcodes are excluded. Orange highlighted in lower pie charts indicates “clonal barcodes,” whose generation probability *P_gen_* are low (< 0.001), enough to be postulated to be introduced in a single PGC. (**C**) Barcode diversity found in individual mice (gray dots). (**D**) The contribution of individual clonal barcodes (*P_gen_* < 0.001) in all recombined barcodes recovered from each mouse. In (C) and (D), values only between neighboring time points were statistically evaluated. Horizontal bars indicate medians. *** P < 0.001, * P < 0.05, two-sided Mann-Whitney-Wilcoxon test (MWW). The transient increase in clone size variation at 3W might reflect the first round of spermatogenesis, which is still ongoing at this stage^44^.

Using this experimental setting, we first analyzed the clonal dynamics of the initially established PGCs (Figure 1A). Following consecutive 4-OHT injections at E6.5 and E6.75 (hereafter collectively referred to as E6.5), tissues including all germ cells, namely the posterior half of the embryo including the allantois at E8.5 and the bilateral testes from E12.5 through 1 year after birth, were collected from multiple male mice. To comprehensively catalogue germline barcodes in these samples, we performed long-read amplicon sequencing of the *Polylox* locus at high sequencing depth (Figures 1A and S3A).

At E8.5, we detected 58.2 ± 13.8 (SD) barcodes per individual (N = 6 mice) (Figures 1B and 1C, and Table S3). Notably, barcode diversity does not directly reflect the number of PGC clones, since some barcodes are more likely to be generated during *Polylox* recombination and are therefore likely to have arisen independently in multiple PGCs^12^. By contrast, barcodes labeling a unique clone (clonal barcodes) will have been generated in a single PGC only. Therefore, by simulating barcode generation as random sequences of *Polylox* recombination^12^, we estimated that the recovered barcode repertoire at E8.5 corresponded to 126.2 ± 35.2 clones (Figure S3B and Supplementary Text). Interestingly, the number of recovered barcodes was halved to 24.5 ± 7.6 at E12.5 (left and right testes combined, N = 13 mice) (Figure 1B and 1C, and Table S4). Thereafter, the barcode count remained stable from late embryonic development through one year after birth. This initial decline highlights a marked reduction in clonal diversity, in which E6.5-labeled PGC clones undergo substantial “pruning” during migration to the embryonic gonads. Once settled in the testes, such PGC clones persist through germline development into adulthood.

Next, we analyzed the PGC clone size. First, we verified that the barcode read frequency is proportional to the number of cells carrying each barcode (Figures S2B and S3C, and Tables S3 and S5). We then focused on clonal barcodes, whose calculated small generation probability (*P_gen_* < 0.001) suggests their generation within a single PGC. We found that the contribution of each clonal barcode recovered at E8.5 was largely even, around 1% of all barcode reads (Figure 1D), consistent with the observation that a single mouse embryo contains 116 ± 4.4 PGCs at this stage^16^. However, by E12.5, when ∼3,500 PGCs are found in bilateral testes (Figure S1F), the distribution of clonal sizes became uneven, ranging from less than 1% (< 35 cells per clone) to about 10% of the total barcode reads (i.e., about 350 cells per clone) (Figures 1D and S3D, and Table S5). While the total germ cell number increased substantially after E12.5, the uneven clonal contributions remained largely stable into adulthood.

Approximately half of clonal barcodes were shared between bilateral testes at E12.5 and beyond, while the remaining half were restricted to one side (Figures S3E and S3F). Clonal barcodes observed bilaterally showed greater read frequencies, indicating larger clone sizes, than those detected unilaterally (Figure S3G). These results indicate that the variation in clone size arises during PGC migration, before they split to colonize the bilateral testes.

### Clone size-dependent transmission to the offspring

To assess how the uneven contributions of E6.5 PGC-derived clones are propagated through SSCs, spermatogenesis, and into the next generation, four E6.5-barcoded males were mated with multiple females from 2 to 10 months of age (Figure 2A). At one year, the males were sacrificed, and barcodes were analyzed in the FACS-sorted SSC pool and in whole testes, representing bulk spermatogenesis (Table S4). We also analyzed the barcodes transmitted to offspring. Since the fathers were heterozygous for *Polylox*, approximately half of the >200 offspring sired by each male inherited barcodes. Sanger sequencing of the offspring’s *Polylox* loci revealed ∼20 barcodes transmitted from each father (Table S6).

**Figure 2.**
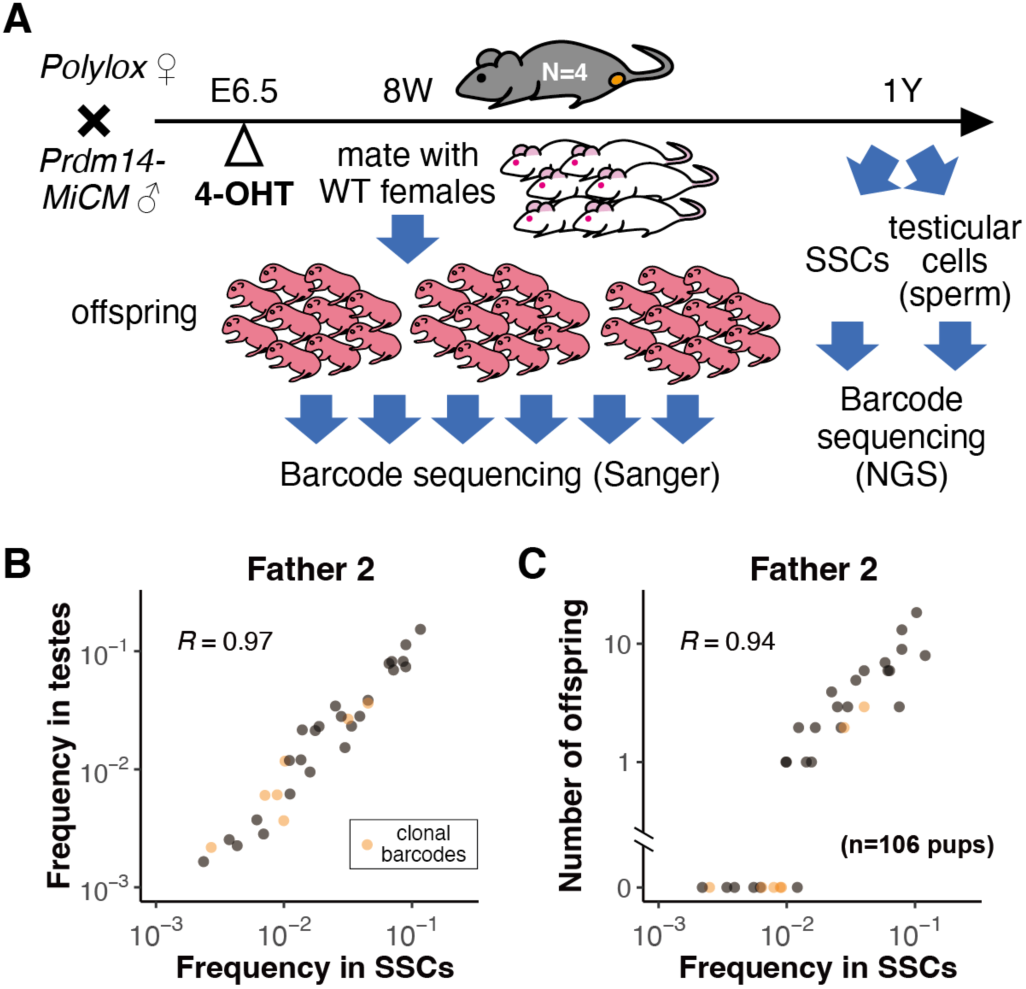
Barcode transmission from SSCs through spermatogenesis to offspring. (**A**) Experimental overview. Four *Prdm14-MiCM*;*Polylox* male mice barcoded at E6.5 were repeatedly mated with multiple females to give rise to a substantial number of offspring till 1Y old. Barcodes transmitted into offspring were determined individually by Sanger sequencing. Then, the fathers were sacrificed and the barcodes included in their SSCs (specifically, undifferentiated spermatogonia) and bulk testicular cells (representing active spermatogenesis) were analyzed by deep-sequencing. Given the heterozygosity of the fathers, approximately half embryos inherit the barcodes. (**B** and **C**) Correlation between barcode compositions in SSCs and the bulk testes (B), and between father’s SSCs and the 106 offspring that harbored barcodes (C). Representative data obtained from one male mouse (Father 2) are indicated; see Figure S4 for those from the other fathers. For SSCs and bulk testis, barcodes from each of the bilateral testes were summed with a 1:1 ratio. “Clonal barcodes” (*P_gen_*<0.001) are highlighted in orange. In (C), actual number of offspring with barcodes are indicated. Spearman correlation coefficients are indicated.

We found that clone compositions showed strong concordance between SSCs, the whole testis (as a measure of sperm), and the offspring (Figures 2B–2C and S4A–S4E). Barcodes with major contributions to SSCs and spermatogenesis were frequently found in offspring, whereas those with minor contributions appeared only infrequently or never. Together, the clonal compositions of E6.5-PGC derived SSCs in fathers are faithfully mirrored in spermatogenesis and transmitted to the next generation.

We then asked whether the PGC clones’ contribution to spermatogenesis and transmission to the next generation shifts over the course of paternal aging (Figure S4F). However, the barcodes observed in offspring showed no signature of paternal age-dependence (Figures S4G and S4H). Therefore, once the SSC pool is established, early PGC-derived clones contribute stably and proportionally to spermatogenesis throughout the reproductive lifespan. This stable and proportional reflection indicates that, although PGC clone sizes (i.e., SSC numbers) differ markedly, clones are functionally equivalent on a per-SSC basis in generating transmissible sperm. Taken together, the observed pruning during embryonic development and subsequent lifelong preservation, and inheritance, of clonal divergence suggest that the early dynamics of PGCs shape their contribution to the next generation while thereafter, reproductive success depends linearly on clone size.

### Stochastic dynamics of PGC clones and prediction of non-arriving PGCs

Motivated by the decisive impact of early clone skewing and pruning on long-term spermatogenesis and transgenerational inheritance, we mathematically analyzed the experimental data. We first asked whether the observed unequal expansion of PGC clones supports selection of fitter PGCs or neutral drift of equally fit PGCs, considering three scenarios: (i) PGC fate is predetermined for survival or loss, (ii) PGCs undergo stochastic expansion dynamics but have heritable fitness differences, or (iii) PGCs are functionally uniform, and neutral drift shapes clonal expansion. These scenarios generate different clone size distributions that we compared with the experimental data.

Remarkably, we found that the measured clone size distributions, at all time points in the fathers as well as in the offspring, reduced to an invariant distribution when rescaling clone sizes by the mean clone size (Figures S5A and S5B). This invariant cumulative distribution was exponential, *e*^−*x*^, where *x* denotes the rescaled clone size, which is the hallmark of neutral drift. This finding excludes scenario (i), and robustly supports scenario (iii), specifically a neutral drift birth-death process in which cells undergo stochastically timed division (“birth”) and loss (“death”) (Supplementary Text). Moreover, we found that a non-neutral birth-death process, with heritable fitness differences between PGCs (scenario (ii)), will yield a clone-size distribution that deviates from exponentiality by developing a heavy tail (Figures S5C and S5D), which is not observed in the measured data. Together, these analyses indicate that the uneven expansion of PGC clones in early development is governed by neutral drift.

We next analyzed the size dynamics of PGC clones across formation, proliferation, and migration to bilateral gonads. To this end, we modeled four plausible schemes based on stochastic birth-death processes of PGCs, and explicitly simulated the generation of *Polylox* barcodes in PGCs through multiple recombination events. Using Bayesian inference, we then evaluated the predictive capacity of each model against the experimental data, including the identity and composition of measured barcodes, the degree of barcode sharing between bilateral testes, and the overall kinetics of PGC numbers between E6.5 and E12.5^16,17^ (Figures S6A and S6B, and Supplementary Text). The model in which PGCs all survive during migration but a fraction of them do not reach the testes explained the experimental data best (Model 1; Figures 3A–3B and S6A–S6F). The cell loss during migration was modeled as a stochastic transition to a “non-arriving PGC” fraction, in which cells are still present and detectable at least until E9.5 but will not colonize the testis. None of the other three models (Models 2–4) explained the measurements with realistic parameters and were therefore discarded (Figure S6A). Thus, Model 1 explains the measured data predicts a non-arriving PGC fraction.

**Figure 3.**
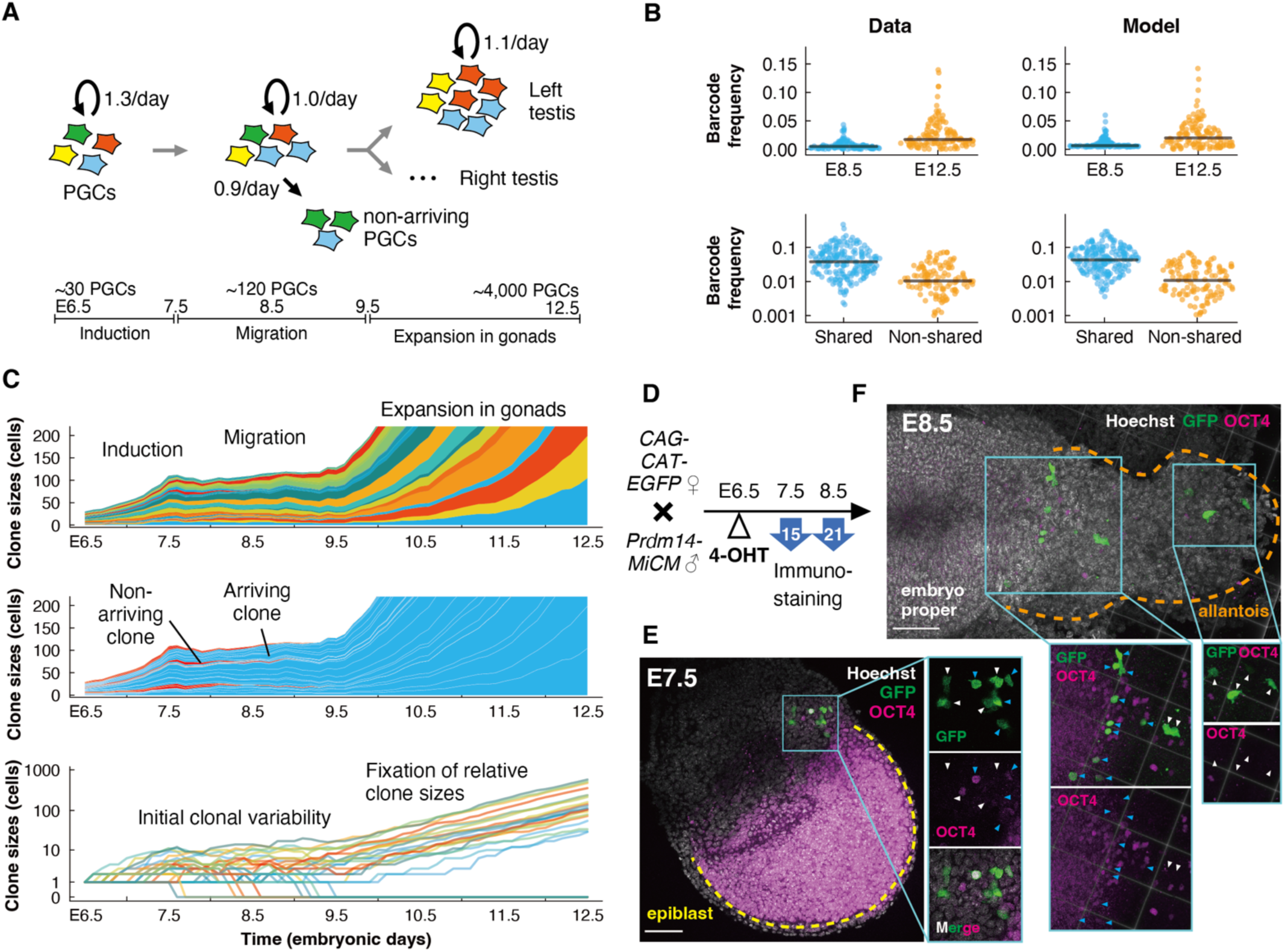
Inference of clonal dynamics of PGCs from E6.5 to E12.5. (**A**) Scheme of the “top model” with parameters inferred from E6.5 PGC-barcoding data. In this scheme, the initial ∼30 PGC clones proliferate through sequential phases of neutral birth-death processes, reaching the left and right gonads at expanded clone sizes. During the migration phase (E7.5–9.5) a non-arriving fraction of PGCs emerges that lost the ability to reach the gonads. The class of birth-death processes was informed by rescaled clone size distributions (Figure S5; Supplementary Text). Different models were tested and fitted on barcode, cell number and sharing statistics, among which this top model was found to be best-fit (Figure S6A). (**B**) Predictive capacity of the top model (right) for a dataset not used for fitting (left). The frequency of clonal *Polylox* barcodes among all recombined barcodes for both measurement time points (top panels; model shows a representative simulation). Barcode frequencies (all except unrecombined) at E12.5 distinguishing barcodes shared and non-shared between bilateral testes (bottom panels). Horizontal lines indicate medians. (**C**) Model prediction of the clonal dynamics of PGCs. Model inference was conducted based on *Polylox* barcodes, followed by re-simulations for individual clones originating from a single cell at E6.5. Clone size dynamics of the re-simulated ∼30 PGC clones were shown in different colors (top panel), arriving (blue) and non-arriving (red) clones (middle panel). Clone size evolution plotted on log-scale (bottom panel), indicating that early stochastic fluctuation in clone size become stabilized by ∼E11, after which clones expanded proportionally. (**D**) Experimental overview for (E) and (F). Following induction by 4-OHT administration at E6.5, GFP-labeled cells in *Prdm14-MiCM*;*CAG-CAT-EGFP* embryos were analyzed at E7.5 and E8.5 for their distribution and marker protein expression. (**E** and **F**) E7.5 (E) and E8.5 (F) embryos, immunostained for GFP (green) and OCT4 (magenta) proteins, cyan and white arrowheads indicating GFP^+^OCT4^+^ and GFP^+^OCT4^−^ cells, respectively. Epiblast (E) and allantois (F) are outlined. (F) 3D visualized images (see Methods). Scale bar, 100μm.

To infer their key quantitative properties, we simulated the clone dynamics of PGCs using Model 1 (Figure 3C). We found that during the early migration phase, when each PGC clone consists of only a few cells, stochastically timed cell division and loss introduced random variations in clone size, estimated to range from one to >12 cells per clone at ∼E9.5. This early heterogeneity in clone size remained fixed during the subsequent expansion phase, as the influence of random division and loss diminishes with increasing clone size (Figure 3C bottom). We inferred that, from the initial ∼30 PGCs at E6.5, ∼20 PGCs contribute to the definitive germline, of which ∼71% contribute bilaterally and ∼29% unilaterally to the testes (Figures S6G and S6H). The ∼10 largest PGC clones make up for ∼84% of germ cells at E12.5 and later times, and eventually the offspring (Figures S6I and S6J). In addition, larger clones at E9.5 are expected more to contribute bilaterally and to remain large till E12.5 (Figures S6H and S6K), consistent with the observed relationship between clone size and bilaterality (Figure S3G). Together, the data-driven model suggests that the variability in PGC clone size is rooted in both neutral drift and loss of PGCs during early embryonic migration, and hence these processes jointly determine subsequent reproductive success.

### Identification and characterization of the predicted non-arriving PGC fraction

Our analysis predicts that a significant fraction of PGCs become non-arriving, failing to settle in the testis and hence not contributing to the definitive germline. We addressed this prediction experimentally. To characterize the cells that had expressed *Prdm14* at E6.5 in vivo, we labeled them with GFP using *CAG-CAT-EGFP* reporter^18^ instead of *Polylox,* to allow for microscopy readout (Figure 3D and Table S7). Following 4-OHT injection at E6.5, we observed GFP-labeled cells exclusively within an extraembryonic region adjacent to the posterior tip of the epiblast at E7.5 (Figure 3E), consistent with the reported PRDM14 expression at this stage^15^. We found that some GFP-positive cells were negative for OCT4, another marker of PGCs (Figures 3E and S7A). Notably, at E8.5, ∼52% (125/239) of GFP^+^ cells were OCT4^+^ and located anteriorly near the hindgut, while the other ∼48% (114/239) were OCT4^−^ and located posteriorly on the allantois (Figures 3F and S7A). Within the GFP^+^OCT4^−^ cells, ∼69% (36/52) expressed extraembryonic mesoderm marker FOXF1^19^ (Figure S7B).

These experiments suggest that about half of the descendants of *Prdm14*^+^ presumptive PGCs lost the germline potential by E8.5 and adopted a different fate. This is consistent with the view that PGCs commit to the germline only after colonizing the gonads, under the influence of gonadal signals and DAZL expression^20–22^. In line with this, recent single-cell analyses also indicate that *Prdm14*^+^ cells constitute common precursors of PGCs and extraembryonic mesoderm, whose PGC potential becomes progressively restricted as their BMP responsiveness increases^23^. Together, these complementary findings converge on a coherent view that a subset of *Prdm14*^+^ cells establish the definitive germline by activating a core set of germline-identity genes (including *Prdm14*, *Oct4*, *Blimp1*, *Tcfap2c*, and *Dppa3*)^3,24,25^, as they migrate into the embryonic gonads. During this process, some cells may migrate improperly^26^, being eventually eliminated by apoptosis^27,28^. Hence, the observed clonal pruning and non-arriving PGCs may reflect the multistep nature of germline development.

### Turnover of gonocyte sub-clones within parental PGC clones

After colonization in the testis, the composition of E6.5-labeled PGC clones remained stable from the late embryogenesis through postnatal and adult stages, with individual clone sizes increasing in parallel with the overall germ cell population^29^. However, while these “parental” clones persisted, it remained unknown whether the constituent sub-clones were proportionally maintained or instead subject to loss and expansion.

To address this question, we barcoded the gonocytes at E11.5 and analyzed the composition of recovered barcodes at subsequent time points (Figure 4A and Table S8). The total barcode number decreased from E12.5 (416 ± 171 per individual) to 1 week after birth, around which SSCs are established (242 ± 50) (Figure 4B). Following this period, the number of recovered barcodes remained stable until 1 year old. Focusing on clonal barcodes (*P_gen_* < 3 × 10^−5^; indicating single gonocyte-derived sub-clones), we estimated that the fraction of sub-clones surviving from embryonic to adult stages is ∼50% (Figure S8A). Further, at E12.5 and E13.5, contributions of individual clonal barcodes were relatively even around 0.03%, which became diverged by 1 week after birth and maintained until one year old (Figure 4C). Rescaled clone size distributions are consistent with neutral drift dynamics (Figure S8B).

**Figure 4.**
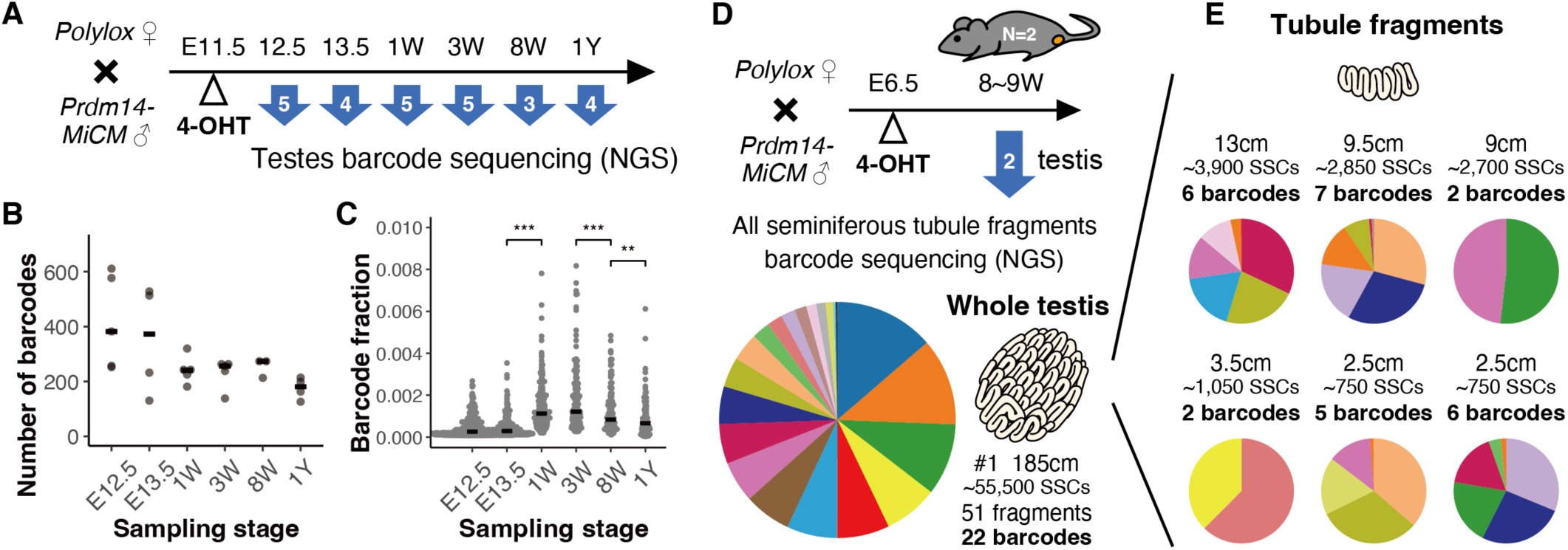
Barcode analysis after late induction and in fragments of seminiferous tubules. (**A**) Experimental overview. *Prdm14-MiCM*;*Polylox* male embryos were barcoded at E11.5, whose barcodes in germline were analyzed at the indicated time points from individuals of indicated numbers shown in arrows. (**B**) Barcode diversity found in individual mice. Horizontal bars indicate medians. No significant changes were detected between nearby stages (two-sided MWW). (**C**) The contribution of individual clonal barcodes (*P_gen_* < 3 × 10^−5^) in all recombined barcodes recovered from each mouse. *** P < 0.001, ** P < 0.01, two-sided MWW. Horizontal bars in (B) and (C) indicate medians. (**D**) Experimental overview is shown in top. Testis from each of the two adult *Prdm14-MiCM*;*Polylox* mice (8∼9W old) induced at E6.5 were decapsulated to collect all the fragments of seminiferous tubules. Then, barcodes in all the dozens of collected fragments were read individually, compiled to reconstruct their composition over an entire testis, and summarized in a pie chart (see Method for detail). (**E**) Examples of long and short tubule fragments obtained in (D), with barcode reads therein. In (D) and (E), same color among pie charts indicates same barcode. See Figure S9 for full dataset for this and the second testis. Estimated SSC numbers in each fragment shown (see Methods).

These findings indicate that, in contrast to the high stability of the PGC clones, their sub-clones undergo loss and expansion during the late embryonic and perinatal periods. Interestingly, although the E11.5-labeled sub-clones became stable during adulthood, it has been revealed that adult SSC-derived clones (i.e., “sub-sub-clones”) also undergo loss and replacement during homeostasis^30,31^. This highlights a hierarchical mode of clonal dynamics, in which stabilization occurs at large scale, while continual turnover persists at small scale.

### Segregated spatial distribution of PGC clones in seminiferous tubules

Thus, we asked how the PGC clone dynamics stabilize in later stages despite ongoing subclonal turnover, and whether stabilization is linked to the drastic changes in tissue environment. After dispersed migration through embryonic tissues, PGCs colonize the embryonic testes and become partitioned among ∼11 distinct testis cords by E13.5 (Figure S1G). Testis cords develop into seminiferous tubules through postnatal lumenization, ultimately extending up to 2 meters in total length per testis in adults^4,32,33^. Through this process, germ cells are confined within a distinct, quasi-one-dimensional tubular geometry, which may restrict clonal intermixing.

To gain insight into the relevance of this compartmentalization, we examined the spatial distribution of germline clones within seminiferous tubules. Following barcoding at E6.5, all seminiferous tubules from two testes, derived from two adult male mice, were collected in fragments ranging from 1 to 13 cm in length (Figure 4D, and Tables S9 and S10). Strikingly, tubule fragments, whether long (∼10 cm) or short (< 4 cm), contained only a minor subset of the collective barcode repertoire, typically 2–7 barcodes (Figures 4E and S9A–S9C). Furthermore, sub-fragments cut from longer fragments tended to share a common set of barcodes (Figure S9D). These findings indicate limited clonal intermixing within seminiferous tubules.

### Modeling the germline clone behavior through development and homeostasis

Then, to quantitatively capture how PGC clones behave within seminiferous tubules, we extended our previous model (till E12.5) into adulthood, by incorporating the tubular anatomical configuration (Figures 5A and S10). In this scheme, at E12.5, germ cells were segregated and aligned within a one-dimensional arrangement of ∼11 seminiferous tubules, which elongated unevenly^32^ until 2 weeks after birth when Sertoli cells ceased proliferation^34,35^ (Figure S10D and Table S2). SSC population is assumed to expand in proportion to the tubule elongation till this stage and then remain constant. We also incorporated the observed ∼50% loss of the E11.5-labeled PGC sub-clones (Figure S8A) and the previously reported stochastic neighbor turnover of SSCs during homeostasis, in which loss of a SSC is coupled with the duplication of one of the neighboring SSCs^31^ (Supplementary Text).

**Figure 5.**
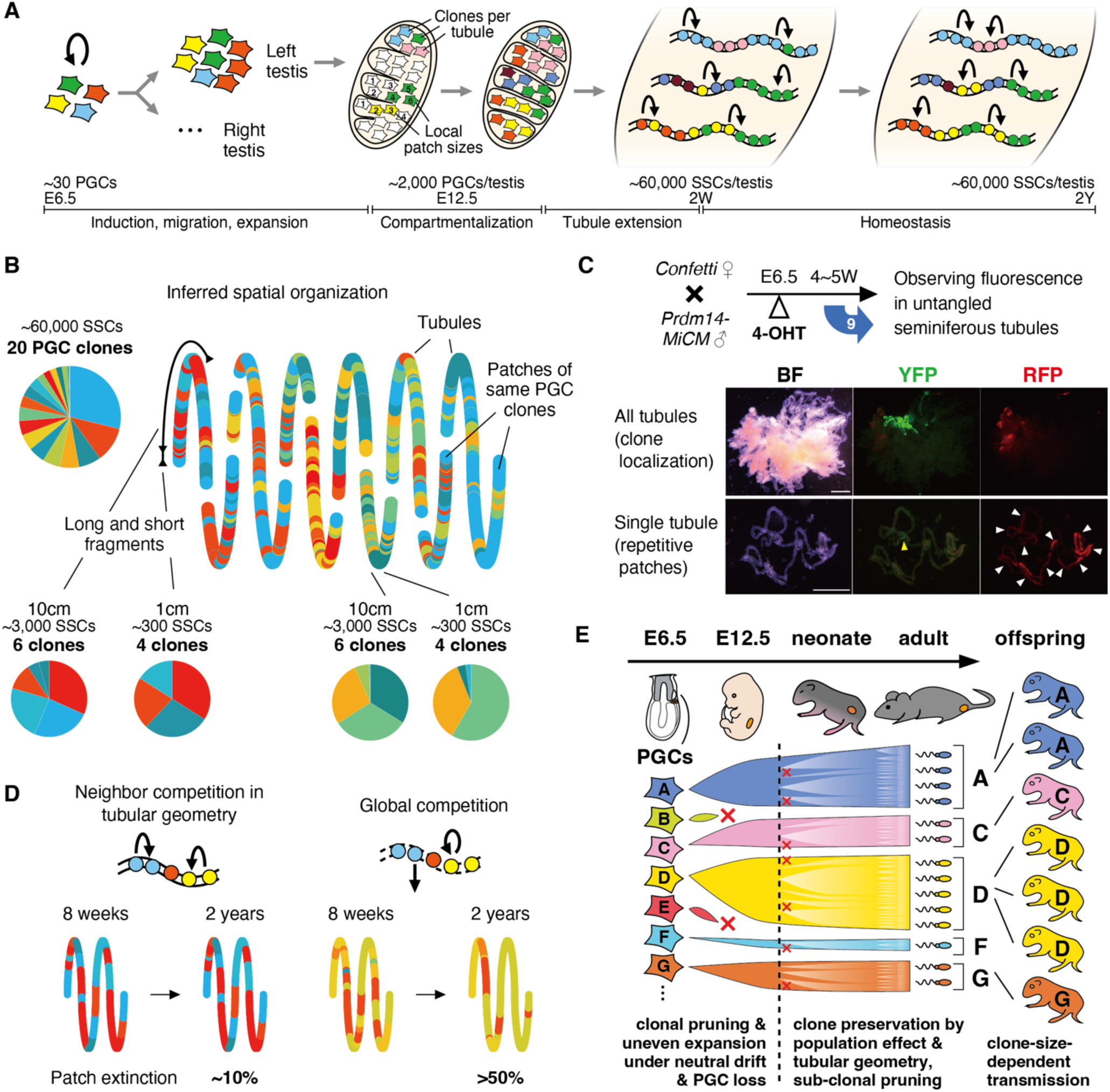
Modeling the lifetime clonal dynamics of male germline. (**A**) Scheme for modeling the lifetime dynamics of the ∼30 initial PGC clones at E6.5. Following the modeling over early phases till E12.5 (Figure 3), additional models were developed to capture the germline fate behavior over extended times, including the compartmentalization into newly formed testis cords (E12.5), seminiferous tubule extension till 2W, and persistent homeostasis from 2W to 2Y of age. See Supplementary Text in detail. (**B**) A typical example of the inferred spatial organization of E6.5 PGC clones at 8 weeks after birth, simulated using the sequential model summarized in (A). Within a single testis containing ∼60,000 SSCs, 20 PGC clones survived and were distributed across 11 separate tubule loops. These clones were not evenly distributed; rather, each tubule loop harbored only a subset of clones. The clones formed a mixed, recurrent patchwork structure in which short segments contained a disproportionately large number of PGC clones (e.g., 4 clones in a 1-cm segment), compared to long segments (e.g., 6 clones in a 10-cm segment), consistent with experimental observations (Figures 4D–4E and S9). (**C**) Visualization of PGC clone distribution using the *Confetti* multi-fluorescence lineage reporter. Experimental design (top). Fluorescence images (middle and bottom) of gently untangled whole seminiferous tubules from a single testis of a *Prdm14-MiCM;Confetti* mouse labeled at E6.5 (2-3m Right), harboring large YFP⁺ and RFP⁺ patches (middle), or a dissected tubule fragment (2-3m tubule 4) containing a small YFP⁺ patch and a cluster of non-contiguous RFP⁺ patches, indicated by yellow and white arrowheads, respectively (bottom). Scale bar, 2 mm. (**D**) Model prediction of the lifetime dynamics of local patches under different modes of adult homeostatic regulation. In the neighbor competition model (left), which incorporates the geometry of seminiferous tubules, local patch lengths remain highly stable with a minimal patch extinction over 2 years life span. In contrast, a hypothetical global competition model with the same differentiation output but agnostic of spatial geometry (right) shows a pronounced disintegration of the local patch structure. (**E**) A schematic diagram summarizing the clonal dynamics of mouse male germline from development to the next generation.

In this modeling scheme, we found that the spatial arrangement of the clones in later stages was strongly influenced by their distribution upon their segregation into testis cord compartments at E12.5. To guide our experiments, we modeled computationally a range of plausible spatial arrangements—from complete mixing to various local patch structures—and then compared the simulated and observed barcode distributions in long and short fragments of adult seminiferous tubules (Figures 4E and S9A–S9C). The observed limited barcode diversity in long fragments indicates that individual PGC clones showed restricted contribution to only a subset of the eleven seminiferous tubule loops. However, intriguingly, a simple contiguous allocation of each clone within a single tubule does not explain the fact that comparable numbers of barcodes are included in short and long fragments. Instead, our modeling infers a locally recurrent patch structure, where small patches derived from a single clone containing ∼150 SSCs on average, alternate along the tubule. Simulations using this structure capture all key experimental observations (Figures 5B and S10A– S10B and S10D–S10E).

To experimentally assess the model-predicted clone distribution patterns, we visualized E6.5 PGC clones 4-5 weeks after birth, using *Confetti*, a multi-color fluorescent protein reporter^36^, instead of *Polylox* (Figure 5C top). With very low recombination efficiencies, colored germ cells were detected in the testes of 5 out of 9 mice, 3 of which showed only a single color (Figure S11A), suggesting that cells marked by the same color belong to a single, or very few, PGC clones. In these samples, cells of a particular color formed clustered patches separated by unlabeled cells, confined to restricted regions within a limited subset of seminiferous tubules (Figures 5C and S11B–S11C). This spatial pattern aligns well with the model prediction, supporting a locally recurrent clonal patch structure achieved during testis development.

### Tubular geometry and population effects stabilize the germline clonal dynamics

In our model reflecting the seminiferous tubule geometry, germ cell clones are stably maintained over extended periods. Extrapolation to 2 years predicted only minimal changes in clone numbers and patch-length distributions (Figures S12A and S12B). Therefore, we considered how such stabilization occurs within seminiferous tubule compartments. First, a *population effect* is likely to contribute: clone size itself buffers size fluctuations and thereby creates stability, with early small clones readily going extinct, whereas later enlarged clones persist. In addition, a *geometric effect* may arise from the one-dimensional arrangement: When clones occupy contiguous stretches, only rare loss-replacement events that occur at boundaries between adjacent clones will influence the clone size dynamics, while the majority of turnover that occurs intraclonally, within a clonal patch, will not.

We evaluated the role of the geometric effect by comparing simulated clone dynamics in models with different geometries. In addition to a truly one-dimensional (1D) model, we implemented a quasi-one-dimensional (q-1D) model^31^ reflecting the cylindrical architecture of tubules, and a global competition model, in which any SSC can compensate for loss of another SSC regardless of their relative positions (Figures 5D and S12C). Interestingly, these models predicted that E6.5 PGC-derived clones stably persist, showing only very little difference in their number and size distribution (Figures S12D and S12E). We reason that the surviving PGC clones had enlarged enough to be refractory to extinction due to the population effect before compartmentalization. In contrast, E11.5-gonocyte-derived sub-clones showed high stability in 1D and q-1D models because these clones originated within these geometric restrictions (Figure S12F). Consistently, the local mosaic patch structure was strongly preserved in 1D and q-1D, but not in the global competition model (Figures 5D and S12G–S12I). Moreover, our 1D model is consistent with enhanced long-term diversity of previously characterized SSC-derived germ cell clones, which originate after the system reached homeostasis (“sub-sub-clones” in the context of this study) within seminiferous tubules^31,37^ (Figure S12J). Importantly, our 1D model also predicts that when an SSC were to emerge with a selective advantage in cell competition (e.g., through a mutation), the expansion of its clone would nevertheless be strongly restricted due to the 1D geometry (Figure S12K). Taken together, these model analyses suggest that the stable persistence of germ cell clones is achieved through the combination of population and geometric effects.

## Discussion

By integrating *Polylox* barcoding with mathematical inference, we quantitatively reconstructed clonal dynamics across the entire lifespan of the mouse male germline, from embryonic development to the next generation. Our analyses reveal biphasic dynamics of PGC-derived clones: an early phase of clonal pruning, during which many lineages are lost, followed by a later phase of long-term preservation, providing an unprecedented bird’s-eye view of male germline development (Figure 5E).

Importantly, the relative sizes, and hence the probabilities of reproductive success, of PGC clones are shaped during early development. This phase of clonal pruning may be critical for establishing the definitive germline, in which cells that have progressed through sequential developmental processes survive and reach the embryonic testes. Our stochastic model accounts for the observed clonal pruning and uneven expansion, yet the molecular mechanisms that underpin the loss of PGCs remain an open question. One possible scenario is that migration of sparsely distributed PGCs through complex embryonic tissues, where extracellular signaling molecules are heterogeneously distributed^26^, introduces stochastic variability in PGC survival and expansion, thereby shaping uneven clonal dynamics.

Once the definitive germline is established, its clonal landscape is largely preserved throughout the male reproductive lifespan. Our model analysis suggests that the observed clone-size variability arises from stochastic fate decisions, rather than fixed intrinsic differences among clones. This clonal landscape is stabilized by the quasi-one-dimensional geometry of the seminiferous tubules. Although SSC clones undergo stochastic loss and replacement^31,37^, clone competition is limited to two interfaces per clone along the tubular axis, markedly slowing clone turnover. Notably, this geometric constraint remains vital even for SSC clones with a selective advantage.

In this context, so-called selfish mutations, which confer a selective advantage to SSCs, can drive non-neutral clonal expansions, leading to a paternal-age-associated increase in the risk of congenital disorders in offspring^1,38,39^. However, such selfish clones remain rare even in aged men, comprising at most on the order of 10^−4^ of total spermatogenesis^1^. By contrast, in two-dimensional epithelia or spatially less constrained hematopoietic tissues, clonal dynamics are far more pronounced than in spermatogenesis^40–43^. In hematopoiesis, for example, mutant stem cell clones that gain a selective advantage can expand to a substantial fraction, reaching clone sizes of 4% or higher^43^. We propose that such tissue-to-tissue differences, including the limited expansion of advantageous SSC clones in seminiferous tubules, are, at least in part, attributable to tissue geometry. Interestingly, these differences resonate with the distinct missions of soma and germline. Somatic cells preserve tissue integrity in both structure and function, and because they are not continuous across generations, they can tolerate some mutational load. The germline, by contrast, while dispensable for the father’s own survival, is responsible for preserving genome integrity for successive generations.

In conclusion, our study reveals how germline fate dynamics shape a diverse clonal output through which genomic and epigenomic information is transmitted across generations. These findings provide a framework for future investigations into how stochastic and selective forces influence fertility, heritable disorders, and ultimately evolution.

## Acknowledgements

We thank the members of the NIBB Yoshida lab, the DKFZ Höfer lab, A. Goriely (Oxford University), and Honda N. (Nagoya University) for discussion, NIBB Model Organisms Facility for animal care, NIBB Trans-Omics Facility for Sequel sequencing, T. Feyerabend for a *Rosa26^Polylox^* mouse strain, M. Saitou, K. Hayashi and G. Nagamatsu for a *Dppa3-ECFP* mouse strain, M. Saitou, H. Ohta and K. Kurimoto for a BAC clone, K. Busch and M. Reth for a *Mer-iCre-Mer* construct, and X. Wang, J. Rößler and D. Postrach for scripts to determine *Polylox* barcodes.

## Author contributions

T.I. performed most of the experiments, with the support of C.K. and A.M.; T.I., M.L., and T.Ni. performed barcode calculations and statistical evaluations; M.L. designed and performed mathematical modeling of clonal/barcode dynamics, with input from T.Ni., N.B.B. and T.H.; Y.K. developed an algorithm to count PGCs; K.Ya. and S.Sh. performed SMRT sequencing and initial data mining; M.L. and K.Yo. performed time-series mathematical analysis of barcode appearance in offspring; T.I., S.Su. and T.Na. conducted cell sorting; S.M. and S.T. established a *Prdm14-MiCM* by microinjection; T.I. and S.Y. designed the experiments; T.I., M.L., T.H., and S.Y. wrote the manuscript with input from all contributors; S.Y., H.-R.R., and T.H. conceived and supervised the study.

## Data and materials availability

All barcode CCS data are archived in DDBJ Sequence Read Archive (DRA) and will be released upon publication. All Sanger sequence reads are also available upon request after publication. The image datasets are available at SSBD:repository upon publication. All code for mathematical modeling was written in the Julia programming language and is available on GitHub. The complete executing code for clonal PGC dynamics is contained in GermlineDynamics (https://github.com/hoefer-lab/GermlineDynamics). Code therein calls methods for *Polylox* barcode simulation and inference from the package PolyloxMechanics.jl (https://github.com/hoefer-lab/PolyloxMechanics.jl). All Bayesian inferences are based on methods from the package ABCdeZ.jl (https://github.com/hoefer-lab/ABCdeZ.jl). During the review process ‘GermlineDynamics’ and ‘PolyloxMechanics.jl’ (as of now private repositories) are available at ***** (Password: *****). The custom programs used for counting the number of PGCs, and all the other calculations for plotting written as R markdown documents, are available upon request. The *Prdm14-MiCM* mouse strain will be available to qualified researchers upon execution of a material transfer agreement (MTA).

## List of Supplementary materials

Materials and Methods

Supplementary Text

Figures S1 to S12

Tables S1 to S10

Additional References *45–58*

## Materials and Methods

### Mice

*Rosa26^Polylox12^*, *CAG-CAT-EGFP*^18^, *Dppa3-ECFP*^45^, and *Rosa26^Confetti36^* alleles were previously described; *Prdm14-MiCM* allele was newly generated in this study, as detailed in the following section. All these mouse lines were maintained in a C57BL/6J background (CLEA Japan or Japan SLC). Males aged 7–52 weeks were crossed with females aged 7–24 weeks to obtain embryos or offspring. The *Prdm14-MiCM* transgenic mice were generated with approval by the Institutional Animal Care and Use Committee (IACUC) of the University of Tsukuba (18-077). All other animal experiments were conducted in accordance with the guidelines for animal experiments of the National Institutes of Natural Sciences (NINS) and approved by the IACUC of NINS (17A107, 18A043, 19A059, 20A063, 21A074, 22A062, 23A064, 24A058, 25A047).

### Generation of a *Prdm14-MiCM* transgenic allele

A bacterial artificial chromosome (BAC) carrying a genome fragment of C57BL/6J mouse spanning the entire *Prdm14* coding sequence plus 157kb flanking sequence (clone RP24-341H14)^15,46^ was modified in *E. coli* (Figure S1A). Briefly, exons 2 to 5 of the *Prdm14* gene were replaced by a *Mer-iCre-Mer*^13,14^ sequence, polyA signal and the FRT-flanked Neo cassette, using RED recombination system^47^. The loxP and lox511 sites on the vector were replaced with *Amp^r^* and *Zeo^r^*genes, respectively. Finally, the Neo cassette was deleted by a transient flippase activation. The resultant, *Prdm14-MiCM* BAC was purified, linearized by PI-SceI digestion, and injected into pronuclei of C57BL/6J mouse zygotes to generate transgenic animals. For G_0_ animals, the presence of transgene was examined by PCR using Taq DNA polymerase (BioAcademia) for 5min at 94°C; (20s at 94°C, 20s at 68–61°C, 7s at 72°C) 8 times with 1°C touchdown; (20s at 94°C, 20s at 60°C, 7s at 72°C) 27 times; 14s at 72°C, with primers *MiCM_3UTR_5For* (5’-GCTTATCGACTAATCAGCCATAC –3’) and *Prdm14_3arm_MidRev* (5’-CCACCTCACTAAGTTGCTGC-3’), amplifying 336bp sequence of the transgene. For F1 and F2 generations, the transgene copy number was quantified by qPCR using THUNDERBIRD SYBR qPCR Mix (Toyobo) and LightCycler 96 (Roche), for 95°C at 60s; (15s at 95°C, 60s at 40°C) 45 times, with primers *Prdm14_qPCR_5For* (5’-CGTTTCGCTAAACTCCTTGG –3’) and *Prdm14_qPCR_5Rev* (5’-TGCAGCCGAGATAGGAGAAG –3’) annealing Intron 1 of *Prdm14*, and *b-actin For* (5’-ACCACAGGCATTGTGATGG –3’) and *b-actin_Rev* (5’-TCGGTCAGGATCTTCATGAGG –3’) as a reference. A transgenic allele with the highest copy numbers at a single locus (line #14) was selected and maintained homozygously (Figure S1B).

### 4-hydroxytamoxifen administration to pregnant mothers

A stock solution of 4-hydroxytamoxifen (4-OHT; Sigma-Aldrich) was prepared by dissolving 25mg 4-OHT, as well as 12.5mg progesterone (Sigma-Aldrich) to sustain pregnancy, into 250μl EtOH and then 2250μl peanut oil (Sigma-Aldrich), using a sonication water bath, and stored at –30℃ until use. Male and female mice homozygously carrying the *Prdm14-MiCM* and the *Polylox* alleles, respectively, were crossed to avoid leaky recombination in zygotes due to maternally-derived MiCM protein accumulated in oocytes during oogenesis. After overnight pair housing, a vaginal plug was checked the following morning, with midday on the day of plug detection designated as E0.5. To induce at E6.5&6.75, two successive doses of 150 µl stock solution (containing 1.5 mg 4-OHT) were orally administered at a 6h interval. For E11.5 induction, a dose of 100μl of stock solution (containing 1mg 4-OHT) was administered orally. For a postnatal sampling of the testes, a Caesarean section was performed at E19.5, and the pups were bred by foster mothers (ICR).

### PGC count

To count the number of PGCs (or gonocytes/prospermatogonia) with and without barcoding, homozygous *Polylox* females were crossed with males carrying *Prdm14-MiCM* and *Dppa3-ECFP* alleles, generating embryos heterozygous for *Polylox*, *Prdm14-MiCM,* and *Dppa3-ECFP*. Testes of these embryos, with 4-OHT administration at E6.5&E6.75 or E11.5, or without administration were harvested at E12.5 and fixed in 4% PFA in PBS for 2h at 4℃. Testes without 4-OHT administration were also sampled at E13.5. The samples were washed with PBS containing 0.1% TritonX-100 (PBST), blocked with 4% Donkey serum in PBST at room temperature (RT) for 3h, incubated with primary antibody solution [rabbit anti-DDX4 (Abcam, ab13840; RRID:AB_443012; 0.05%) supplemented with 4% donkey serum in PBST] at 4℃ overnight, washed with PBST at RT for >1h four times, and incubated in 2^nd^ antibody solution [donkey anti-Rabbit IgG (H+L) Cy3 (Jackson ImmunoResearch, 711-165-152; RRID:AB_2307443; 0.1%) and goat anti-GFP (SICGEN, AB0020-200; RRID:AB_2333100) conjugated with DyLight™ 488 NHS Ester (Thermo Fisher Scientific) (0.1%), supplemented with 4% donkey serum in PBST] at 4℃ overnight. After washing with 1mL PBST at RT for >1h four times, the testes were mounted with Scale SQ^48^, and serial images were acquired for 3D reconstruction using TCS-SP8 (Leica). To count the number of PGCs automatically, captured images were analyzed by custom program written for R software environment (www.r-project.org) with EBImage packages^49^. In brief, PGC masks were created using ECFP and immunostained DDX4 signals by applying Gaussian blur filter followed by local thresholding and size selection. Information of each single PGC across several slices was integrated. PGCs in individual testis cords, obtained from two independent E13.5 testis images, were also counted (Figure S10D upper right).

### General procedure for barcode sequencing, decoding, and filtering

The *Polylox* locus was PCR-amplified from genomic DNA samples using primers #2,427 and #2,450^12,50^ and KOD FX Neo polymerase (Toyobo) with a thermal cycler MiniAmp Plus (Thermo Fisher Scientific) under a thermal cycling condition of initial denaturation (94°C for 2 min) followed by 28 or 30 cycles of 98°C for 10 sec, 64°C for 30 sec, and 68°C for 150 sec, and a final extension (68°C for 7 min). The amplicons were purified using AMPure XP (Beckman Coulter) at a 1:1 ratio, and proceeded for library preparation using a SMRTbell^®^ Express Template Prep Kit 2.0 (Pacific Biosciences) according to the manufacturer’s protocol (101-791-700), and long-read Single Molecule, Real-Time (SMRT) sequencing using Sequel I and Sequel IIe sequencers (Pacific Biosciences) with standard parameters. The obtained circular consensus sequencing (CCS) reads were converted to barcodes, followed by classifying into ‘legitimate’ barcodes (those which are theoretically possible to occur from the original *Polylox* sequence) and ‘illegitimate’ barcodes (those which are expected not to occur), as described previously^12,50^. Given the rarity of illegitimate barcode detection and their distinct read number distribution from the vast majority of legitimate barcodes, we reason that these anomalous barcodes likely arose from PCR or bioinformatics processing errors, rather than from genuine recombination events involving the *Polylox* locus within germ cell genomes. This suggests that legitimate barcodes may include some artefacts, so we filtered out rarely observed barcodes altogether so that the illegitimate barcode reads became less than 5% of total reads (Figure S3A).

### Specificity verification of barcode introduction into PGCs

Triple transgenic embryos carrying *Polylox*, *Prdm14-MiCM* and *Dppa3-ECFP* alleles resulting from the crossing described previously were barcoded by 4-OHT administration at E6.5&6.75 or E11.5. For each schedule, gonads collected from eight embryos at E12.5 were pooled and dissociated by a Liberase solution [0.5 mg/mL DNase (Worthington), 0.26 U/mL Liberase^TM^ TM (Roche), and 18U /mL Britase (Wako) in PBS]. A total of 7,500–8,800 PGCs and ∼300,000 gonadal somatic cells were separately collected as the ECFP^+^ and ECFP^−^ fractions, respectively, using a MA900 cell sorter with Software v3.2.2 (Sony) (Figure S2A), and proceeded for genome PCR amplification (30 cycles) and barcode analysis as described above. We found that recombined barcodes were detected only in the ECFP^+^ fraction sorted from the testes of embryos obtained from 4-OHT injected mothers, with no recombination detected at all in the ECFP^−^ fraction or samples without 4-OHT injection, verifying that barcode recombination occurred in a high germ cell-specific and tamoxifen-dependent manner (Figure S1E).

### Barcode analysis of developing and adult germ cells

To observe the clonal dynamics during development, embryos carrying *Polylox* and *Prdm14-MiCM* alleles were barcoded at E6.5&6.75 or E11.5, followed by harvesting posterior-half embryos at E8.5 (for E6.5&6.75-labeled embryos only) or bilateral testes at time points spanning from E12.5 to 1 year of age. Samples harvested at embryonic stages, i.e., posterior-half embryos and testes after removing the mesonephros, were lysed separately in 16μl protease solution; all the template DNA was directly used for PCR amplification (28 cycles) and barcode analyses. Postnatally, right and left testes were independently collected, removed off the tunica albuginea, lysed in 200–1000μl protease solution followed by DNA purification, and 100– 1000 ng template DNA (representing 1.7 × 10^4–105^ cells) was used for barcode PCR amplification (28–30 cycles) and barcode analysis. To calculate the barcode compositions per individual, barcode reads from bilateral testes were combined based on the assumption that the germ cell numbers were equal between testes.

We found that ∼4% of these double-transgenic mice harbored recombined barcode sequences broadly over somatic cells, likely due to leaky and 4-OHT-independent recombination during very early development. Accordingly, barcode sequencing was performed on samples taken from individuals whose somatic tissues (head or tail, respectively, for samples taken at E8.5 or other stages) were confirmed by PCR amplification to have full-length *Polylox* sequences only. Further, the sex of the E8.5 and E12.5 embryos were determined by PCR amplification of the *Rbm31x/y* genes^51^, and mice after E13.5 were judged by the testis morphology.

Barcode sequencing of individual E13.5 PGCs (Figure S3D), isolated by FACS-sorting of the Dppa3-ECFP+ population (Figure S2A), was performed similarly to that of undifferentiated spermatogonia, as described below.

### Verification that the barcode reads linearly reflect the cell composition

Considering the possibility of biased PCR-amplification, library construction and sequencing, we verified if and how much the composition of the barcode reads mirrors that of cells carrying heterogeneous barcodes. Testes from adult mice barcoded at E6.5&6.75 were dissociated by a Liberase solution (see above), and were incubated in antibody solution [Alexa Fluor 647-conjugated rat anti-CD9 (BioLegend, 124810; RRID:AB_2076037; 0.2%, PECy7-conjugated rat anti-KIT (BioLegend, 105814; RRID:AB_313223; 0.5%) and Brilliant Violet 421-conjugated rat anti-CD324 (BioLegend, 147319; RRID:AB_2750483; 0.1%) in PBS] on ice for 30 min. Then, a CDH1^+^/CD9^+^/KIT^−^ undifferentiated spermatogonia fraction was collected by a MA900 cell sorter with Software v3.2.2 (Sony), as a bulk of 17,000–25,000 cells and in parallel as a separate assembly of hundreds of individual cells (Figure S2B). Barcode analyses through PCR-amplification and amplicon long-read sequencing were conducted as described previously. In parallel, barcodes are also decoded separately in individual cells by Sanger sequencing (Fasmac) of the *Polylox* sequence amplified by nested PCR. The primers used are #2,450 and #494 for the first round, and #2,426 and #2,427 for the second round, respectively^12^. Both rounds of PCR were performed using KOD FX Neo at 25μl scale and the following protocol: 2 min at 94°C followed by 25 rounds of (10s at 98°C, 30s at 64°C, 2min30sec at 68°C) and 7min at 68°C. 1μl of the 1^st^ PCR product was used as a template for the 2^nd^ PCR. The products of the second PCR were treated with ExoSAP-IT Express (Thermo Fisher Scientific) and were sequenced with primers #2,427, #2,450, #2,623 (5′-AGAAGTTGCATACACAGTATTG-3′) or #2,622 (′-ATCATCGAGCTCGTCAACAATG-3′)^12^. A custom script (SangerSeqSingleFilesAnalysis.m from Daniel Postrach) was used to determine barcode elements in sequencing reads.

We found a strong linear correlation between the frequency of barcode reads obtained from the bulk population using deep sequencing and the number of cells whose barcode was individually determined (Figure S3C). We concluded that the barcode frequencies obtained from bulk Sequel deep sequencing faithfully reflect the composition of cell numbers with these barcodes within the population of interest and that the technical biases, if any, are negligible.

### Comparison of barcode distributions between the SSC pool, spermatogenesis, and offspring

Four male mice barcoded at E6.5&6.75 were each crossed with multiple ICR females from 8–9 weeks old. Because the males heterozygously harbored the *Polylox* transgene, about half of the offspring inherited barcodes via sperm. The fathers were sacrificed at 1 year of age, whose right and left testes were separately dissociated using a Liberase solution (see above). Half of the dissociated cells were used to FACS-sort the “SSCs” fraction (i.e., CDH1^+^CD9^+^KIT^−^ undifferentiated spermatogonia; 12,000–20,000 SSCs per testis) as described above (Figure S2B), while the other half were directly proceeded as “spermatogenesis” fraction without sorting, for amplicon sequencing as described previously to gain barcode reads. In parallel, the offspring’s tail tips were digested by Cica Genius AN (Kanto Chemical) to prepare the DNA, followed by PCR-amplification of the *Polylox* sequence using KOD FX Neo with the temperature condition of 2 min at 94°C followed by 32 rounds of (10s at 98°C, 30s at 64°C, 2min30sec at 68°C) and 7min at 68°C, using primers #2,427 and #2,450. PCR products were treated with ExoSAP-IT Express, and barcodes were sequenced in the same way as single-cell-derived barcodes (see above). Temporal bias of barcode appearance in offspring was evaluated using the methods detailed in the Supplementary Text.

### Visualization of *Prdm14^+^* cell clones using a fluorescence reporter

To visualize the spatial distribution of the descendants of cells expressing *Prdm14* at E6.5&6.75, homozygous *CAG-CAT-EGFP* females were crossed with homozygous *Prdm14-MiCM* males to conceive embryos carrying *Prdm14-MiCM* and *CAG-CAT-EGFP* alleles heterozygously. Following 4-OHT administration to pregnant mothers at E6.5&6.75, whole embryos and posterior-half embryos were harvested at E7.5 and E8.5, respectively, and fixed in 4% PFA in PBS for 2h at 4℃. After washing with PBST, the samples were blocked with 4% Donkey serum in PBST containing 0.1% Hoechst 33342 (Thermo Fisher Scientific) and incubated for 3h at RT. Then, the blocking solution was switched to the primary antibody solution [anti-GFP (goat, Sicgen, AB0020-200; RRID:AB_2333100; rat, Nacalai Tesque, 04404-84; RRID:AB_10013361; or chicken, Aves Labs, GFP-1010; RRID:AB_2307313), together with rabbit anti-OCT4 (Abcam, ab181557; RRID:AB_2687916), and with or without goat anti-FoxF1 (R&D Systems, AF4798; RRID:AB_2105588); all used at 0.1%, and supplemented with 4% donkey serum in PBST], and the samples were incubated at 4℃ overnight (>16h). Afterwards, samples were washed with PBST at RT for >1h four times, and incubated in the secondary antibody solution [any one of anti-Goat IgG (H+L) Alexa 488 (705-545-147; RRID:AB_2336933), anti-Rat IgG (H+L) Alexa 488 (712-545-153; RRID:AB_2340684) or anti-Chicken IgY (H+L) Alexa 488 (703-545-155; RRID:AB_2340375), together with anti-Rabbit IgG (H+L) Cy3 (711-165-152; RRID:AB_2307443), and with or without anti-Goat IgG (H+L) Alexa 647 (705-605-147; RRID:AB_2340437); all donkey-derived (Jackson ImmunoResearch), used in 0.1%, and supplemented with 4% donkey serum in PBST] at 4℃ overnight. After washing with PBST at RT for >1h four times, the samples were mounted using Scale SQ^48^, and serial images were acquired for 3D reconstruction with a TCS-SP8 confocal microscope (Leica). Three-dimensional visualization was performed using ClearVolume^52^. To assess potential antibody cross-reactivity, we omitted each primary antibody individually from the antibody mix and confirmed the specific loss of its corresponding signal. The sex of E8.5 embryos was determined as described above, and no sex-dependent differences were observed.

### Barcode analysis of seminiferous tubule fragments

To address the spatial distribution of barcoded clones over seminiferous tubules, the testes of two E6.5&6.75-barcoded male mice were obtained at 8–9 weeks of age. For each mouse, one of the testes was lysed for barcode amplicon sequencing as described earlier to verify the degree of recombination; the other was extensively disentangled to collect virtually the entire seminiferous tubule length in as long fragments as possible. Fragments 4cm or longer were cut into sub-fragments. Genomic DNA purified from each of these (sub)fragments was used for amplicon-sequencing to determine the barcode composition included therein. For each fragment, barcodes with <1% read frequency were filtered out because they were likely due to contaminated germ cells from other fragments during sampling. To estimate the length of tubule segments occupied by cells with each barcode, its read frequency was multiplied by the length of the fragments from which the barcode was yielded. The number of SSCs in each fragment was estimated by multiplying its length by 30 cells per millimeter^53^.

### Multi-color reporter observation of PGC clones in seminiferous tubules

Homozygous *Rosa26^Confetti^* females were crossed with homozygous *Prdm14-MiCM* males to obtain fetus that obtain both genes heterozygously. After 4-OHT administration at E6.5&6.75 and Caesarean section as in the same way with barcoding procedure, seminiferous tubules of bilateral testes obtained from males between 29 and 36 days old were untangled thoroughly and fixed in 4% PFA in PBS for 1h at 4°C.

Followed by washing with PBST, the fluorescent images of seminiferous tubules were obtained using a M165FC stereoscopic microscope (Leica) with GFP-LP, RFP-LP and ET-CFP filters. Green signals were regarded as cYFP but not nGFP according to their localization in the cytoplasm. We could not find an ECFP signal, possibly because the mECFP cassette was not activated by recombination or the mECFP signal was not strong enough to be distinguished from background signals. Because we found clear cYFP and cRFP signals in only limited areas in 5 out of 9 mice (Figure S11A), we reasoned that recombination occurred infrequently, and labeled cells derived from only one or a few PGCs at E6.5&E6.75.

### Statistics

No statistical methods were used to determine the sample size. Mice under identical experimental conditions were randomly sampled at different developmental time points, and in principle all dissected samples with successful tissue processing were included. This resulted in datasets in which each time point was represented by samples derived from more than two mice from more than two independent litters. For the developmental time-course barcode analysis, two E12.5 embryos from the same litter with barcode induction at E6.5&6.75 exhibited unexpectedly high barcode diversity, exceeding the expected number of PGCs at the time of induction, and were therefore excluded as outliers relative to the other 13 embryos. Except for these two embryos, all time-course samples showed consistent tendencies in barcode distribution. For postnatal analyses, only mice that survived Caesarean section and were successfully fostered were included. Blinding was not applied because the study assessed temporal changes within the same labeling condition. As the analyses did not involve comparisons between different experimental conditions, the risk of bias was minimal. Because barcode distributions were not expected to be normal, two-sided Mann-Whitney-Wilcoxon tests were applied to assess differences between adjacent time points, and Spearman’s rank correlation coefficients were calculated to evaluate the relationship between barcodes in two samples.

### Mathematical modeling of clonal PGC dynamics

Methods and theory for the modeling of clonal PGC dynamics are detailed in Supplementary Text. In brief, we modeled cellular dynamics of division, migration, loss and spatial organization following clones of the emerging E6.5 PGCs. The cellular dynamics of PGC clones were simulated alongside *Polylox* barcode recombination, to infer key biological insights from multiple aspects of the experimental barcoding data. Namely, the reduction of clonal diversity in the embryo, followed by the maintenance of the survivors, and the spatial organization of these clones in the testes over the murine life span. We investigated mechanism for the postnatal stability of PGC clones, incorporating the specific geometry of the seminiferous tubules in the testes shaping competition of the SSC clones emerging from the PGCs.

## Supplementary Text

The methods and theory for modeling germline clonal dynamics are detailed in the following sections *Supplementary Methods* and *2. Supplementary Equations*.

### 1. Supplementary Methods

#### Scaling of clone size distributions

We assessed whether clone sizes from E6.5&6.75 PGC barcoding conform with a neutral birth-death process; the theoretical basis for this procedure is given in Supplementary Equations a. We first applied the read threshold to filter spurious barcodes and subtracted the threshold from the kept clone sizes, so that the smallest possible clone size is again at one read (this mitigates censoring effects). Afterwards extant clone sizes were rescaled (divided) by their mean clone size, and the complementary cumulative probability distribution of the rescaled clone sizes was compared to the theoretical scaling law (Figures S5A and S5B). Further, this scaling of neutral clonal competition was compared to processes with inherent and persistent clonal biases (i.e., selective advantages) among the PGC clones at E6.5, confirming neutral stochasticity as the principal source underlying the observed clone size variation (Figures S5C and S5D, and Supplementary Equations b).

#### Mechanics of *Polylox* recombination

*Polylox* barcode recombination was modeled as a Markov process in continuous time, each cell being subjected to a recombination rate *r*. We assumed this rate is the same for all cells during the entire recombination window; particularly, independent of the current barcode state. *Polylox* recombination was then simulated stochastically by starting all cells with the unrecombined barcode ‘123456789’, selecting the next cell and time to recombine, conducting a recombination (that is, changing the cell’s barcode state) and continuing this procedure until the end of the recombination window. The new barcode state was sampled from a 1,866,890×1,866,890 transition matrix specifying all legitimate *Polylox* recombinations^12^. Matrix values were assumed to be uniform, apart from a bias *w*_exc_ ∈ (0, 1) that tunes chances of excisions vs. inversions. For example, barcode ‘123456789’ has 45 possible recombinations, 25 inversions (preserving barcode length 9) with probabilities *c* (1 − *w*_exc_) each and 20 excisions (leading to lengths 7, 5, 3 or 1) with probabilities *c w*_exc_each, and *c* = 1⁄(25 (1 − *w*_exc_) + 20 *w*_exc_). Where applicable, such barcode recombination was tied to other reactions of cellular dynamics (cell division, loss, etc.) in a single model (see subsequent sections). At divisions, both daughter cells inherited the barcode of the mother.

#### Modeling PGC dynamics from E6.5 to E12.5

To infer the lifetime clonal dynamics of the ∼30 PGCs induced at E6.5, we set up model inferences drawing from multiple barcoding experiments, starting with analysis stages E8.5 and E12.5. We reasoned that the accurate evolution of clonal loss and clone sizes are revealed from such a time-series inference, in contrast to single snapshots alone. Further, we designed the approach to be capable to switch between *Polylox* barcodes (introduced in a more continuous time window from E6.5 to ∼E8.0) and clones derived from single E6.5 PGCs.

First, we created four models of clonal PGC dynamics spanning from E6.5 to E12.5 as continuous time Markov jump process with sequential phases of neutral birth-death processes (as tested in section ‘Scaling of clone size distributions’). All models were initialized with thirty PGC clones at E6.5 that proliferated as a pure birth process with division rate λ_1_ until E7.5, and continued to divide during a subsequent migration phase with rate λ_2_until E9.5. At E9.5 cells were randomly distributed to two compartments, representing left and right gonads (later testes), where they proliferated with rates λ_3_ until E12.5. The models differ in their potential PGC loss during the migration phase (Figure S6A). In model 1, PGCs were lost with a rate μ to a non-arriving PGC fraction that was not distributed to the gonads at E9.5.

Second, we added *Polylox* barcodes to these models. Each cell was equipped with an internal barcode state, starting with the unrecombined barcode ‘123456789’ in all cells at E6.5. Alongside the cellular dynamics, cells internally recombined their *Polylox* barcode with a rate *r* and excision bias *w*_exc_ from E6.5 to E8.0 (see section ‘Mechanics of *Polylox* recombination’); after E8.0, barcodes were considered stable. Hence, given values for its biological parameters (λ_1_, λ_2_, λ_3_ and μ), each model simulated specific E6.5 PGC dynamics. And, given additional values for the recombination parameters (*r* and *w*_exc_), these PGC dynamics were simultaneously probed with *Polylox* barcodes, keeping track of barcode identities for all cells during the simulation.

These models were applied to *Polylox* barcoding data, using Bayesian parameter inference and model selection from ABCdeZ.jl. We compared data and models via summary statistics, iterating over many different parameter combinations. Flat priors were chosen: pure birth rates λ_1_and λ_3_uniform in [0.5, 1.75] (per day), the migration parameters λ_2_ and μ uniform in [0, 1.5] (per day), the recombination rate *r* uniform in [0, 3] (per day) and the excision bias *w*_exc_uniform in (0, 1). Summary statistics for the data were computed as median over all 6 and 13 mice for E8.5 and E12.5 analyses, respectively. A first set of summary statistics informed about clonal diversity, using distributions of *Polylox* barcode number (all barcodes used, no *P_gen_* cutoff) over barcode lengths (9, 7, 5, 3, 1) and minimal recombinations (0, 1, …, 11), and sharing of barcodes between left and right gonads (E12.5 only). A second set of summary statistics informed about total PGC cell numbers^16,17^, using read distributions over barcode lengths and minimal recombinations rescaled to 200 (E8.5) and 2000 cells (E12.5 for left and right each). For each parameter set, models were simulated once until E8.5 and once until E12.5, producing analogue model summary statistics from simulated *Polylox* barcodes. The arriving and non-arriving PGC fractions were merged for the E8.5 readout if applicable (models 1 and 2). After fitting each model to the data, posterior model probabilities were obtained, and within each model, posterior parameter distributions.

To extrapolate to the clonal dynamics of the initial ∼30 PGCs, the top model (model 1) was re-simulated with median values of the biological parameter posteriors, keeping track of the founding E6.5 PGC clones (instead of *Polylox* barcodes) for all cells during the simulation.

#### Clone initiating cell numbers for E8.5 analysis

We fitted a recombination-only model to observed barcode numbers of E8.5 analysis after E6.5&6.75 PGC barcoding, to assess whether inferred cell numbers conform with the PGC numbers in the recombination window (E6.5 to ∼E8.0). The barcode data of each of the six mice (E8.5 analysis) was transformed to summary statistics of absolute barcode number distributions over *Polylox* barcode length (9, 7, 5, 3, 1) and minimal recombination number (0, 1, …, 10). For each mouse, the recombination model was fitted to these data, producing the respective summary statistics from simulations of *Polylox* barcode recombination in *N* cells at a recombination rate *r*. Prior distributions were uniform on intervals *N* ∈ {1, …, 500} and *r* ∈ [0, 4] (per day); the recombination window was fixed to 1.5 days and excision bias to 75% (as inferred from the PGC E6.5 to E12.5 dynamics). The median of the posterior distribution for *N* was reported as most likely cell number.

#### Spatial and lifetime model of PGC clonal dynamics

We extended the clonal PGC dynamics inferred during embryonic development (Figure 3, section ‘Modeling PGC dynamics from E6.5 to E12.5’) in two aspects. First, starting around E12.5, PGCs in the gonads localize in cord structures that become the seminiferous tubules of the testes (Figure S1G). We wondered how PGC clones participate in the different tubules and how they are locally distributed inside the effectively one-dimensional tubules. Second, we aimed to model the dynamics of PGC clones over murine lifetimes, both globally and locally. Together, this extended model provides a quantitative view of the spatial and lifetime dynamics of E6.5 PGC clones.

We continued with PGC clone sizes from the left gonad after the lateral split E9.5 (symmetric to right gonad), as inferred from the embryonic model. Clones were expanded again as a pure birth process to reach ∼2000 cells total at E12.5. At E12.5 the expanded clones were then distributed on a one-dimensional spatial lattice, representing the later spatial organization of postnatal tubules: We introduced two parameters *c*_1_and *c*_2_that allow to describe various types of spatial organization, based on a theory for patch formation (Supplementary Equations c). In short, *c*_*i*_ ∈ [0, 1] specify the probabilities that the next cell seeded to the spatial lattice is from the same clone as the previous cell. Cells were distributed on a single lattice according to *c*_1_, the lattice split up into 11 different cords, and, lastly, cells were distributed within each cord lattice according to *c*_2_. Hence, *c*_1_ captures clonal aggregation effects in the gonads before cord formation, leading to biased cord participation, while the product *c*_1_*c*_2_describes the local mixing within a cord after cord compartmentalization, specifying the patch structure as it develops towards postnatal analysis. As corner cases, this procedure contains fully randomized spatial arrangements (*c*_1_ = *c*_2_ = 0), as well as fully aggregated scenarios (*c*_1_ = *c*_2_ = 1) where all clones are laid out one after the other.

The developmental expansion from E12.5 to 2 weeks postnatally was modeled as an expanding birth-death process on the spatial tubule lattices. Single cells on the lattice across all cords were expanded with a mean amplification of 30 (from ∼2000 PGCs to ∼60,000 SSCs per testis^53^) and a single-cell survival chance of 50% (specified by E11.5 barcoding, Figure S8A). We factored in the appropriate level of cord-to-cord heterogeneity in this mean expansion (roughly ranging from 20 to 40, Supplementary Equations d and Figures S10D–S10E). Daughter cells of expanded clones were seeded as immediate neighbors, leading to elongated tubule lattices.

From 2 weeks onwards, clonal dynamics were modeled as a homeostatic process using neighbor competition with an average stem cell replacement time of once every 6 days (following previous studies^31,54^). Concretely, whenever a stem cell differentiated, its empty lattice position was immediately taken by a random left or right neighbor that duplicated in the process. Furthermore, effects of different homeostatic regulations were analyzed (see section ‘Neighbor and global competition during adult homeostasis’).

By using E9.5 clone sizes from the embryonic phase inference and fixing many model parameters as specified above, *c*_1_ and *c*_2_ were the only remaining parameters to be inferred from data. Again, we resorted to simulation-based inference by ABCdeZ.jl. Data of *Polylox* tubule fragments (E6.5&6.75 barcoding, 8– or 9-weeks analysis) were converted to summary statistics of complementary cumulative probability distributions, specifying the fraction of segments exceeding a certain barcode number (for three different categories of segment lengths and two independent mice; Figure S10A). Starting from flat uniform priors over [0, 1] for *c*_1_and *c*_2_, analogue model summary statistics were computed iteratively and compared to the data. For the inference, again *Polylox* barcodes were taken as clone sizes. To obtain model summaries, tubules were simulated until 8 weeks postnatally and then cut into fragments matching fragment lengths of the data (using a conversion of ∼30 SSCs per mm tubule^53^). Fitting the model to the data yielded a correlated posterior distribution for *c*_1_ and *c*_2_.

To gain the spatial and lifetime dynamics of E6.5 PGC clones, we extrapolated again to the initial ∼30 E6.5 PGCs by re-simulating this extended model with *c*_1_and *c*_2_ samples from their two-dimensional posterior and keeping track of PGC clone identity (instead of *Polylox* barcodes). The model was evaluated at various embryonic and postnatal time points, and local patch sizes as well as global clone sizes (summing all patches of same PGC identity) were inspected.

#### Neighbor and global competition during adult homeostasis

We investigated the potential advantages of the specific geometry of seminiferous tubules for the lifetime stability of PGC clones. The standard neighbor competition was simulated in one– or quasi-one-dimensional (1D or q-1D) tubular geometries, and compared to a hypothetical reference model with global competition that is agnostic towards geometric dimensionality or form. Simulations showed that the tubular geometries provide superior stability in contrast to the non-geometric reference; a feature that is further supported by Supplementary Equations e and f.

The spatial and lifetime PGC dynamics were re-simulated as before (see section above), until the postnatal homeostatic phase at 2 weeks was reached. From that point onwards, the tubules were simulated in three different ways (Figure S12C). First, as a neighbor competition process on a 1D lattice where left and right neighbors may replace each other (as before, section ‘Spatial and lifetime model of PGC clonal dynamics’). Second, as neighbor competition on a q-1D cylindrical lattice, where left/right and below/above neighbors may replace each other. The circumference contained 5 cells with periodic boundaries^31^, and hence ∼12,000 units in the longitudinal dimension for a total of ∼60,000 cells. Third, a reference global competition process, modeled as a critical birth-death process where events of cell division and differentiation balance out on the population average. Clones in the birth-death process develop independently of their neighbors and were assumed to be non-mixing at patch boundaries. All three scenarios were simulated with a stem cell division rate λ = 1⁄6 (per day; as before), so that the production of differentiated progeny and sperm are identical.

Simulations tracked E6.5 PGC clones for various time points across the three scenarios. Global clone sizes and local patch sizes were collected to compute end-point-correlations and extinction levels. To quantify the gain of patch stability coming from the inferred average patch size of ∼150 cells (Figure S12B), a hypothetical no-patch case was computed by using the single-cell patch extinction probabilities of a 1D neighbor competition model (Figures S12E and S12H). The single-cell extinction is given by *p*(ℓ_*t*_ = 0) = 1 − *e*^−2λ*t*^(*I*_0_(2λ*t*) + *I*_1_(2λ*t*)) (Supplementary Equations e), and hence the extinction chance for a “patch” of ℓ cells, being instead scattered on the lattice, was computed by *p*(ℓ_*t*_ = 0)^ℓ^. Another set of simulations tracked single cells introduced at E12.5 (PGC sub-clones) in otherwise identical models, to compare clonal diversity over time with the E11.5 barcoding data (Figure S12F). Further, extinction probabilities of single SSC-derived clones (PGC sub-sub-clones) under homeostasis were compared for the 1D neighbor and the global competition model (Figure S12J), setting ℓ_0_ = 1 in Supplementary Equations e.

Finally, the effect of the tubular geometry was analyzed in a scenario where a single E6.5 PGC clone had a selective advantage *s* during the postnatal homeostatic phase. Tubules at 2 weeks were forward simulated with 1D neighbor competition and global competition, increasing the stem cell division rate of the selected clone to λ(1 + *s*) with *s* = 1%, and reading out the spatial configuration at 2 years postnatally (Figure S12K). The tubular geometry with neighbor competition effectively confined the patches of the “mutated” PGC clone, in contrast to exponentially expanding patches under global competition (Supplementary Equations f).

#### Temporal transmission bias from fathers to pups

We assessed whether PGC clones have a temporal bias in their contribution to pups over the father’s life span. To that end, we computed the times between occurrences of the same barcode and aggregated these inter-birth intervals for all barcodes and the four fathers. A value of zero days corresponds to barcode coincidences in two or more pups on the same day. To obtain a model reference of unbiased barcode transmission, the barcodes were randomly shuffled across the pups of a given father and inter-birth intervals were recomputed as done for the data. The data distribution of inter-birth intervals was compared to the unbiased model based on repeated simulations providing median and 95% credible intervals (Figure S4G). Further, the data was reduced to barcodes with a maximum barcode frequency of 10%, 5% or 2% and simulations of the unbiased model were recomputed to obtain right-tailed *p*-values under a null distribution for clonal coincidences^55^ (Figure S4H).

### 2. Supplementary Equations (a, b, c, d, e and f)

#### a, Rescaled clone size distributions of single PGCs

Analysis of rescaled clone size distributions (as done in Figures S5A and S5B) shows that clonal dynamics of PGCs conform with a neutral birth-death process. Due to this observation, we implemented the models for the barcoding data based on these stochastic processes (dynamics from E6.5 to E12.5, Figure 3, and also extended from E6.5 to 2 weeks, Figure 5). The clone sizes from E6.5 barcoding reveal the nature of the clonal dynamics in the expanding embryonic phase. The scaling law then applies even for adult analysis stages, as clones stabilize before entering the adult homeostatic phase (compare with Figure 3) and continued turnover is neutral among surviving clones (see also Supplementary Equations b, Figures S5C and S5D). We briefly outline the well-known theory of birth-death processes, to state the scaling law that we applied to the barcoding data.

In birth-death processes, two types of transitions are possible: An individual cell may divide with probability λ(*t*)d*t* (“birth”) and leave the compartment of interest with probability μ(*t*)d*t* (“death”) in any short time span d*t*. The division and loss rates, λ(*t*) and μ(*t*), are the same for all cells in the considered compartment. Particularly, they are independent of the clone identity and current clone size, and as such the process is neutral. The rates may however vary as a function of time *t*; this includes our models with sequential phases. For this process, the probability distribution of clone sizes *X*_*t*_ = *x* at time *t* for any initial clone size *X*_0_ = *x*_0_ are well-known (simple non-homogenous birth-death process; Bailey, 1964^56^). Focusing on clones derived from single cells, *x*_0_ = 1, the distribution is given by *p*(*x*) = (1 − α) (1 − β) β^*x*−1^ (for *x* ≥ 1) and *p*(0) = α, where α = 1 − 1/(*e*^*p*(*t*)^ + *A*(*t*)) and β = 1 − *e*^ρ(*t*)^/(*e*^ρ(*t*)^ + *A*(*t*)) with 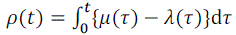 and 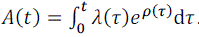. The solution includes the time-homogenous processes (critical, sub– and supercritical). For *Polylox* data it is natural to condition this distribution on survival (as we can only observe extant clones that did not go extinct before analysis). Under this condition, *X*_*t*_ ≥ 1, the distribution simplifies to a shifted Geometric distribution with *p*(*x*) = (1 − β) β^*x*–1^, where *x* ∈ {1, 2, … }, and mean clone size *E*(*X*_*t*_) = 1⁄(1 − β). So, while considering a broad class of processes, all clone sizes fall on the same distributional shape with only one effective parameter—its mean clone size.

Thus, we rescale the extant clones with their mean. We consider *W*_*t*_ = *X*_*t*_⁄*E*(*X*_*t*_) and obtain *p*(*W*_*t*_ > *w*) = β^⌊w⁄(1−β)]^, with floor function ⌊ ⌋. In the continuous limit for expanded clones (β → 1, *w* > 0), the law for the rescaled clone sizes further simplifies to *p*(*W*_*t*_ > *w*) = *e*^−w^. Both expressions were used (Figure S5A and S5B) and pose a simple but strong test of neutrality and compatibility with the class of birth-death processes.

Of note, other neutral processes (e.g., homeostatic Moran process without selection in two or more spatial dimensions) settle on the same scaling law^57^. We further note that non-simultaneous introduction of clonal barcodes *in vivo* should not be a problem for the scaling analysis: Clonal barcodes in model simulations (being created over a 1.5-day recombination window) still scale (Figure S6D), showing that this effect does not distort clone sizes significantly in our system. Lastly, the random partitioning of clones into left and right gonad is equivalent to binomial sampling in each gonad. This intermediate sampling step is already covered within the above birth-death processes, as binomial sampling is mathematically equivalent to a short pure death process, and hence the stated distributions and scaling laws remain unchanged.

#### b, Theory of inherent and persistent clonal bias

In the following we derive a theory to investigate whether, and to which degree, clone sizes that conform with the neutral scaling law (Supplementary Equations a) actually indicate an underlying neutral mechanism. To that end, clonal dynamics are considered where some clones exhibit inherent and persistent selective advantages, termed “clonal bias,” deviating from purely neutral behavior. We find a criterion that directly links non-neutral clone sizes to the fractions of clone size variability that are caused by clonal bias and neutral stochasticity, respectively. This theory confirms that the principal source underlying the observed germline clone size variation is neutral stochasticity (85% – 100%).

Following up on Supplementary Equations a, we start by considering single-cell derived, extant clones driven by birth-death dynamics. Under neutral clonal behavior, clone sizes *X* = *x* at the time of analysis follow the distribution *p*(*x*) = (1 − β) β^*x*–1^, *x* ∈ {1, 2, … }, where β, and hence the mean clone size *E*(*X*) = 1⁄(1 − β), is identical for all clones. To introduce systematic clonal bias and selective advantages, we now lift this assumption and allow for clone-specific dynamics. Concretely, we draw mean clone sizes *g* that may be different across clones, and then sample the observed size *X* for each clone based on the above shifted geometric distribution satisfying the clone-specific mean size *g*. Mathematically, we have *E*(*X* | *g*) = 1⁄A1 − β(*g*)) ≡ *g* for individual clones. This construction implies that each individual clone has underlying division and loss rates identical for all its constituent cells; however, these rates may vary across cells of different clones. To parametrize *g* on the mean and variance level, we set *E*(*g*) = μ and Var(*g*) =^2^. The neutral case is then contained with σ = 0, while σ > 0 implies that some clones will obtain a selective advantage (for events *g* > μ) and others are disadvantaged (*g* < μ). The mean observed clone size across all (potentially biased) clones is calculated to *E*(*X*) =μ, independent of neutral or non-neutral dynamics. In contrast, the variance across all clones is given by Var(*X*) = (μ − 1)μ + 2σ^2^ leading to a coefficient of variation (CV) of 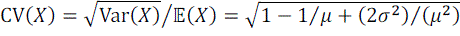. Here, while neutral dynamics approach a maximal CV of one, non-neutral dynamics can generate clone size variation beyond this limit.

To obtain a “scale-free” view on clone size variability, i.e. independent of the mean clone size μ, we introduce a dimensionless bias parameter *b* that quantifies the excess of non-neutral clone size variation beyond the neutral baseline. Concretely, we set 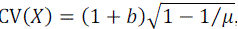, where *b* ≥ 0 and 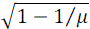 is the CV of the neutral case (σ = 0). From this, one obtains the conversion 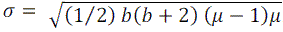. Such clonal bias, quantified via *b*, allows to directly compare the shapes of scale-free clone size distributions (i.e., rescaled) for neutral (*b* = 0) and various non-neutral clonal dynamics (*b* > 0), as done in Figure S5C (*b* ∈ {0.0, 0.05, 0.2, 0.5, 1.0}). To visualize concrete distributions, *g* was sampled from a shifted gamma distribution, setting *g* ∼ Gamma(α, θ) + 1 (with shape α and scale θ) and α = ((μ − 1)⁄σ)^2^ and θ = (μ − 1)⁄α, where σ was specified by the above conversion from *b* and μ. Then *E*(*g*) = μ and Var(*g*) =^2^ as desired.

It remains to be investigated to what extent inherent clonal bias (or selective advantages) actually causally contributes to the observed variation of clone sizes (the shapes of these rescaled distributions). For this, the observed extant clone sizes, *X*, are correlated with their inherently biased means, *g*. If initial biases among clones faithfully predict and determine later “winner” and “loser” clones (i.e., big and small clones), this equates to a high correlation. In contrast, a low correlation reveals that—even if biases among clones may exist—neutral stochasticity dominates effects of clonal bias so that the “winner” and “loser” clones are simply determined by chance. In this sense, the squared Pearson correlation *r*^2^(*X*, *g*) determines the fraction of the clone size variance that is explained by clonal bias (Var(*X*) explained by *g*) and is computed subsequently. First, the joint expectation can be derived to *E*(*Xg*) = *E*(*g*^2^) = μ^2^ + σ^2^. Then, it follows that *r*^2^ = cor(*X*, *g*)^2^ = Var(*g*)⁄Var(*X*) = σ^2^/A(μ − 1)μ + 2σ^2^). Finally, by using the above conversion to express σ in terms of the clonal bias *b* and mean clone size μ, one obtains the result *r*^2^(*X*, *g*) = (1⁄2) A*b*(*b* + 2))/A1 + *b*(*b* + 2)), which becomes independent of μ. Hence, shapes of the rescaled clone size distributions, as set by the clonal biases *b*, directly relate to the fraction of clone size variability, *r*^2^, attributable to inherent and persistent selective advantages. The remaining fraction, 1 − *r*^2^, is hence the clone size variability explained by neutral stochasticity. This result provides a simple yet powerful criterion to assess the contributions of neutral vs. selective clonal dynamics directly from the observed variation of clone sizes.

Comparing the theoretical clone size distributions (Figure S5C) to the experimental distributions (Figures S5A and S5B) reveals no skewness in the data as it would be expected from inherent clonal bias. Additionally, already a bias of 20% (*b* = 0.2) deviates noticeably from the neutral distribution (Figure S5C), and, as such, may be considered the maximal clonal bias hidden in the data. This extent of clonal bias corresponds to a causal contribution of *r*^2^ = 15%, at maximum, to the observed clone size variability, originating from a maximal variation in mean clone sizes of about 47%, as given by 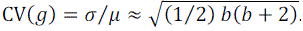. Conversely, at least, 1 − *r*^2^ = 85% of the variability of clone sizes is caused by neutral stochasticity (Figure S5D). Together, these results show that clonal dynamics that appear neutral also indicate an underlying dominantly neutral mechanism. The neutral clone sizes of the germline hence pose a strict bound on the contribution of inheritable and persistent factors to PGC clone sizes.

#### c, Theory of patch formation

We derive a theory for the patches that a clone forms when being distributed on a one-dimensional spatial lattice, representing the forming cords in the embryo and the later tubules in postnatal testes. Patch formation is described by a local correlate of how much a clone identity of a cell predicts the clone identity of its neighboring cells. Concretely, we introduce a probability *c* ∈ [0, 1] that the next cell on the lattice, viewed from left to right, is occupied by a cell of the same clone. This principle, starting from a patch with its first cell, induces patch lengths (or sizes) ℓ ∈ {1,2, … } that follow a shifted geometric distribution (ℓ − 1 “successes,” i.e. further cells of the same clone, before the first “failure”) with probabilities *p*(ℓ) = (1 − *c*) *c*^ℓ–1^. However, for a clone of *N* cells total, a patch length has to be bounded by ℓ ≤ *N*, and the fragmentation of the clone leads to *n* ∈ {1, …, *N*} distinct patches of sizes ℓ_*i*_, such that 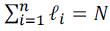. Hence, we aim to derive, conditioned on the clone size *N* and *c* given, the distribution of the number of patches *p*(*n*|*N*) and the distribution of patch lengths *p*(ℓ_*i*_|*n*, *N*) for each patch *i*.

To compute *p*(*n*|*N*) one notes that the inverted probability *p*(*N*|*n*) is given by a shifted negative binomial distribution, as the sum of independently formed patches that are geometrically distributed with *p*(ℓ_*i*_). We have *N*|*n* ∼ NegBin(*n*, 1 − *c*) + *n* with probabilities 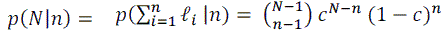. Then Bayes theorem allows the conversion to *p*(*n*|*N*). With an uninformed prior for the number of patches, *p*(*n*) = 1/*N*, one obtains 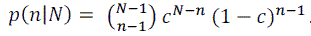. Thus, the number of patches follows a modified binomial distribution, *n*|*N* ∼ Bin(*N* − 1, 1 − *c*) + 1, with a mean patch number *E*(*n*|*N*) = (*N* − 1)(1 − *c*) + 1, and noteworthy corner cases of a single patch containing all *N* cells at once (*n* = 1 if *c* = 1) vs. *N* single-cell patches (*n* = *N* if *c* = 0).

Given *c* and clone size *N*, and with that also the patch number *n* from above, the distribution of patch sizes *p*(ℓ_*i*_|*n*, *N*) is obtained in iterative manner. We introduce ℓ_1_, the size of the first patch and, stochastically independent, 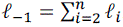, the combined sizes of the patches after the first. It follows *p*(ℓ_1_ = *x*|*n*, *N*) = *p*(ℓ_1_ = *x*) *p*(ℓ_–1_ = *N* − *x*)/*p*(ℓ_1_ + ℓ_–1_ = *N*) (where the right-hand side probabilities are shifted negative binomial distributions from above), leading to 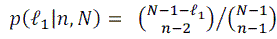, with patch sizes in the range 1 ≤ ℓ_1_ ≤ *N* − *n* + 1. Further, we derive that all subsequent patch sizes can be obtained iteratively: The patch size ℓ_*i*_ is given by the same distribution, only by updating *N*_*ne*w_ = *N* − ℓ_*i*–1_ and *n*_*ne*w_ = *n* − 1; for example, the second patch length is given by 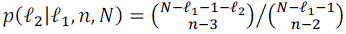. We implemented an algorithm to robustly evaluate the involved ratio of binomial coefficients, termed the BinRatio(*n*, *N*) distribution in code. The mean value of this distribution is E(ℓ|*n*, *N*) = *N*/*n*, while the mean patch length over all (random) patch numbers *n* is computed to E(ℓ|*N*) = (1 − *c*^*N*^)⁄(1 − *c*).

Of note, this theory is closed under repeated patch formations: Fragmenting a clone of *N* cells with *c*_1_ ∈ [0,1] and fragmenting the resulting patches (viewing patch lengths as clone sizes ℓ_*i*_) another time with *c*_2_ ∈ [0,1], is identical to a single fragmentation using the product value *c* = *c*_1_*c*_2_ on the initial clone size *N* directly. This indicates that complex biological fragmentation processes may be well-captured descriptively by a single or few parameters, correlating clone identities of cell neighbors.

Combining above derivations, a list of patch sizes can be generated over all clones, by first sampling the number of patches for a given clone and then sampling the size of each patch iteratively. In summary, this theory provides a full stochastic framework to spatially distribute clones with given clone sizes on a one-dimensional lattice, encapsulated in a single parameter *c* that specifies the statistics of the number and lengths of the formed patches.

#### d, Theory of fetal patch elongation dynamics

We develop a model for the dynamics of patch sizes inside the expanding tubules, spanning from cord formation (E12.5) to the postnatal tubules (2 weeks) at which the homeostatic phase is entered. The model accounts for growth differences in the cords (that is, some cords elongate to long tubules in the adult, some to short), at a magnitude that we validate through independent features in the barcoding data.

Each gonad (the later testis) forms ∼11 cords (the later tubules) that roughly expand from 2,000 PGCs at E12.5 to 60,000 SSCs at two weeks after birth (and thereafter), meaning that PGCs amplify 30-fold on average. We compare the coefficient of variation (CV) of cell numbers at these two time points, to assess to what extent PGC expansion is heterogeneous across different cords. Embryonic cord cell numbers have a CV_E12.5/E13.5_ ≈ 0.31 (Figures S1G and S10D), which increases to CV_adult_ ≈ 0.43 (Nakata et al., 2015^32^; where tubule T6 was excluded as a likely merger two cords; Figure S10D). Subsequently a theory is derived to show that this increase in variation cannot be explained by homogenous stochasticity of PGC expansion but warrants systematic growth differences between elongating tubules.

We assume that a cord *i* has *X*_*i*_ cells initially (E12.5), elongating to *Y*_*i*_ cells postnatally. Within a given cord, single PGCs (labeled by *j*) follow an expanding (supercritical) birth-death process, captured in the random variables *Z*_*i*j_, such that the sum of all single PGC expansions generates the elongated tubule, i.e. we have 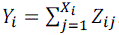. We inform single-cell expansions by the mean E(*Z*_*i*j_) = μ = 30 (from 2,000 to 60,000, over all cords), and survival chance *p*(*Z*_*i*j_ ≥ 1) = 1 − *a* = 50% (E11.5 barcoding, Figure S8A). To probe heterogeneity between single-cell expansions across cords, we allow that E(*Z*_*i*j_|*i*) = *g*_*i*_, where *g*_*i*_ is a (random) per-cord expansion; all cells of the same cord amplify with same mean *g*_*i*_, but cells of different cords may be subject to varying mean expansions. Then, it follows that E(*g*_*i*_) = μ is specified, while Var(*g*_*i*_) = σ^2^ expresses the level of cord expansion variation through the additional parameter σ. The homogeneous case is contained with σ = 0, and σ > 0 encodes heterogeneous expansion. Critically, this framework enables the derivation of means and variances of all involved quantities. We obtain the mean relation E(*Y*_*i*_) = μ E(*X*_*i*_) and a variance of tubule cells after elongation as

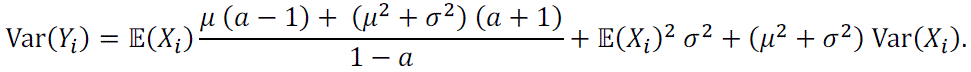

Hence, σ can be computed directly from the embryonic and adult CVs using

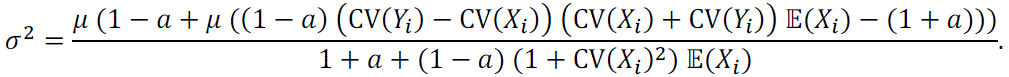

When inserting values of μ = 30, *a* = 0.5, E(*X*_*i*_) = 2000/11, CV(*X*_*i*_) = CV_E12.5/E13.5_ ≈ 0.31 and CV(*Y*_*i*_) = CV_adult_ ≈ 0.43, we obtain σ ≈ 7.5. This indicates substantial variation in cord expansion, with some cords expanding 20-fold, and others 40-fold, to have on average a 30-fold increase in cell numbers. In simulations, we parametrize the mean cord expansion by *g*_*i*_ ∼ Gamma(α, θ) + 1 (with shape α and scale θ) and set α = ((μ − 1)⁄σ)^2^ and θ = (μ − 1)⁄α, satisfying *E*(*g*_*i*_) = μ and Var(*g*_*i*_) = σ^2^ as desired (Figure S10D, for an overall scheme). Thus, solely by comparing cell number variation in embryo and adult, we can specify a model for the expansion of the testis cords becoming elongated tubules at appropriate mean and variance levels.

As the barcoding data is not being used to set up the expansion model or specify σ, we wondered if a feature in that data allows to test the model. If the substantial heterogeneity in the effectively one-dimensional elongations were true, one would expect the following: On average, the barcode patches, created at cord formation, are more stretched out in longer tubules at their final length, while in less elongated tubules they are compressed in a shorter length. Hence, we compared the mean barcode number in small fragments (that came preferentially from shorter tubules) vs. the mean barcode number in sub-fragments that are pieces of a long fragment (preferentially a long tubule); where both sets of (sub)fragments were subsampled to an identical fragment length distribution. Indeed, we observe a slight shift between these mean values in both mice (∼0.3 barcodes in mouse 2), while (without fitting these data) the model predicts a similar shift (∼0.2 for mouse 2; Figure S10E); by contrast, no shifts (for σ = 0) or drastic shifts of more than one barcode (σ ≫ 7.5) are possible in theory but not supported by these data. This independent feature of the barcoding data validates the determined level of cord variation of the expansion model with σ ≈ 7.5.

#### e, Theory of patch length variation in adult homeostasis

We develop a theoretical view of the patch length variation that explains the increased stability of patches in the neighbor competition model (tubular geometry) compared to the global competition model (agnostic of spatial geometry). To that end we derive the time evolution of the coefficient of variation (CV, standard deviation divided by the mean) for typical patch lengths in both models.

In the global competition model birth (cell division) and death (here representing biological cell differentiation) events of stem cells are balanced, on average, on the population level. As a model we used the critical birth-death process, with a birth rate λ equaling the death rate. The mean and variance of patch sizes ℓ_*t*_ over time *t*, starting with an initial size of ℓ_0_ ≥ 1 cells, are known to be E(ℓ_*t*_) = ℓ_0_ and Var(ℓ_*t*_) = 2ℓ_0_λ*t* (Bailey, 1964^56^, Equations 8.54, 8.55, 8.58; Supplementary Equations a) with a probability of extinction as *p*(ℓ_*t*_ = 0) = (λ*t*⁄(1 + λ*t*)) ℓ^0^. Hence, patch lengths under global competition vary as 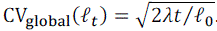.

In the neighbor competition model stem cell differentiation is compensated by the immediate division of a neighboring cell (semantically equivalent to a replacement of a cell by one of its neighbors). The model achieves the same differentiation output as the global competition model when such replacements happen with the same rate λ. Importantly, a patch of cells with the same clone identity maintains an unaltered size when replacements take place within the patch boundaries (cells replacing cells of same clone identity). Hence the evolution of the patch length is only driven by the stem cell dynamics at the patch boundaries. On a one-dimensional lattice, we observe shifts of the left boundary with a rate λ, as the patch increases by one cell when its leftmost cell replaces its left neighbor (rate 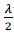), and the patch shrinks by one when its leftmost cell is replaced by its left neighbor of a different clone identity (rate 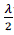). With analogous dynamics at the right boundary, we obtain a total rate of 2λ, independent of the current patch length ℓ_*t*_. Hence one realizes that the patch length ℓ_*t*_ is described by a continuous-time symmetric random walk with an absorbing state at ℓ = 0 for patch extinction. The solution of this process is known as *p*(ℓ_*t*_ = ℓ) = *e*^−2λ*t*^ (*I*_ℓ–ℓ0_ (2λ*t*) − *I*_ℓ+ℓ0_ (2λ*t*)) (for ℓ ≥ 1) and 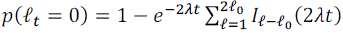, where *I*_v_(*x*) is the modified Bessel function of first kind and ℓ_0_is the patch length at time *t* = 0 (Heathcote and Moyal, 1959^58^). The mean follows as E(ℓ_*t*_) = ℓ_0_ (identical to a random walk without absorbing state). The variance is given by Var(ℓ_*t*_) ≈ 2λ*t* for our typical parameter regime (low probabilities of patch extinction for neighbor regulation, see below). This implies that patch lengths under local neighbor competition vary as 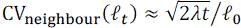.

Critically, the variance under neighbor competition does not scale with the initial patch size, in contrast to the linear dependence under global competition. For typical parameters of *t* = 365 days, λ = 1⁄(6 days) and our inferred mean patch size of ℓ_0_ ≈ 150 cells (Figure S12B), we obtain patch extinction probabilities as *p*(ℓ_*t*_ = 0) ≈ 0.087 (global) and *p*(ℓ_*t*_ = 0) ≈ 1.9 × 10^−38^ (neighbor); and hence, CV_global_ ≈ 0.90 and CV_neighbour_ ≈ 0.07. This explains the drastic differences in long-term patch stability between the two competition models. Of note, we may extrapolate to a human testis by estimating the size of adult patches that reach a similar stability (in terms of the CV) over a comparable human life span. Assuming the same division rate of individual stem cells and considering *t* = 50 years, a CV of 7% is reached at ℓ_0_ ≈ 1050, suggesting a seven times larger human stem cell pool of about 420,000 cells per testis.

#### f, Effect of tubular geometry under selection

We investigate the effect of the tubular geometry by equipping a single PGC clone with a selective advantage *s* > 0 during the postnatal homeostatic phase. This selective advantage may originate from a mutation or epigenetic modification that was acquired during development, and enhances stem cell division in all tubular patches of the selected clone in the adult. The selected clone competes with neutral PGC clones within the testis which follow the dynamics as described in Supplementary Equations e.

Under global competition, the selected clone divides with an increased rate λ(1 + *s*) per cell; hence, its individual patches expand with a mean of E(ℓ_*t*_) = ℓ_0_*e*^λ*st*^, where ℓ_0_ is the patch size at 2 weeks postnatally. Under neighbor competition (in 1D), the selected clone has an increased stem cell replacement rate of λ(1 + *s*). Here, in contrast, the selective advantage is neutralized within each patch, as all neighboring cells share the same advantage. A patch of the selected clone realizes its advantage only at the patch boundaries, gaining cells with a constant rate of λ(1 + *s*) per patch, while losing cells with rate λ. Hence, patches of the selected clone expand as an asymmetric random walk^58^ with a mean of E(ℓ_*t*_) ≈ ℓ_0_ + λ*st*, under negligible patch extinction (Supplementary Equations e).

With a selective advantage of *s* = 2%, a neutral division rate of λ = 1/(6 days) and a time span from 2 weeks to 2 years postnatally, a patch of ℓ_0_ = 150 cells expands to only ∼152 cells on average under neighbour competition, while the patch would amplify >10-fold under global competition, reaching an average size of ∼1632 cells. Thus, neighbor competition within the tubular testis geometry effectively confines selected clones to an extremely slow linear expansion, in contrast to a fast exponential expansion in the hypothetical non-geometric environment.

**Figure S1.**
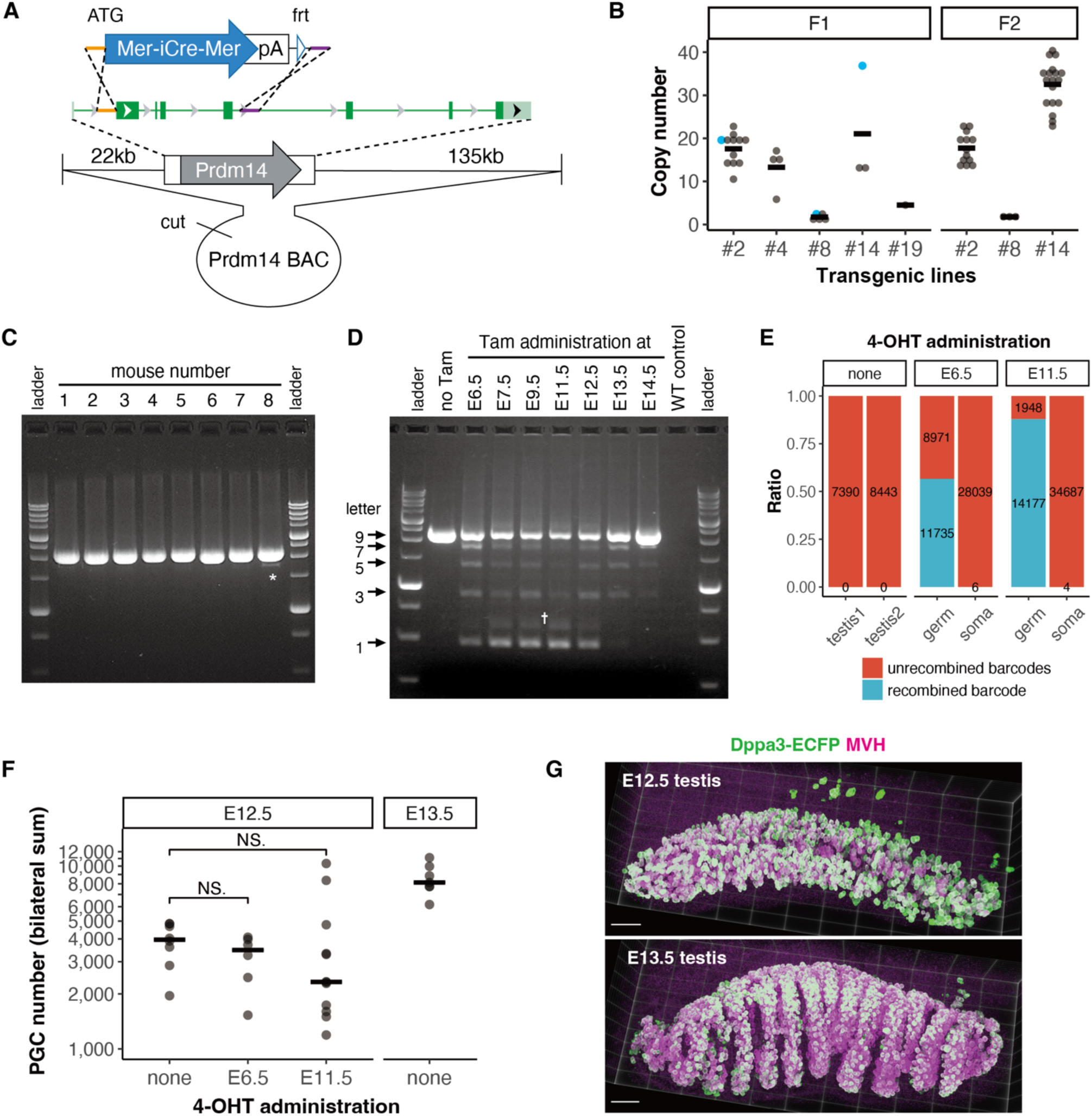
Generation of *Prdm14-MiCM* transgenic alleles. (**A**) Design of the *Prdm14-MiCM* BAC plasmid used for generating the *Prdm14-MiCM* transgenic allele, in which exons 2 to 5 was replaced by *Mer-iCre-Mer* CDS and a polyA (pA) addition signal. (**B**) Copy number of the *Prdm14-MiCM* transgene in independent multiple lines, measured by genomic qPCR. Horizontal bars indicate means. The F_1_ individuals colored in cyan were backcrossed to obtain the F_2_ offspring. Given the largely unchanged copy number between F_1_ and F_2_ generation, these transgenes were highly expected to be integrated in a single locus in lines #2, #8, and #14. (**C**) PCR-amplification of the *Polylox* locus from the tail DNA of *Prdm14-MiCM;Polylox* mice with tamoxifen administration at E6.5, showing predominant amplification of 9-letter length fragments. The lane with asterisk incudes a shorter amplified band (7-letters), likely due to a leaky recombination in early embryos. 3 out of 74 mice (4%) with or without tamoxifen administration showed such recombined short fragment(s) in tail DNA, which were excluded from subsequent analyses. (**D**) Representative examples of the PCR-amplified barcode sequences from testes of 3-weeks-old mice, which have received tamoxifen administration at different developmental time points as indicated. The amplified fragments with five lengths, corresponding to the 9, 7, 5, 3, and 1 letters, indicate that homologous recombination occurred between the loxP sequences contained in the *Polylox* sequence, as expected. Consistent with the expression window of Prdm14^15^, prominent recombination was observed between E6.5 and E13.5. Dagger indicates an unidentified fragment around 600bp, which were informatically excluded from the analysis. Mice that received no tamoxifen showed no evidence of barcode recombination. (**E**) Verification of tamoxifen-dependent and germ cell-specific barcode induction. Testes from 3-weeks-old *Prdm14-MiCM*;*Polylox* mice without 4-OHT administration contained no recombined barcodes (left). The actual numbers of Sequel sequence reads are indicated on each colored bar. *Prdm14-MiCM*;*Polylox;Dppa3*-*ECFP* triple transgenic embryos were induced by 4-OHT administration to mothers at E6.5 or E11.5 (middle and right). Then, germ cells (ECFP^+^) and gonadal somatic cells (ECFP^−^) were FACS-sorted from pools of gonadal cell suspension prepared from eight E12.5 embryos obtained from a single litter (2 males and 6 females for E6.5; 3 males and 5 females for E11.5). Recombined barcodes were detected exclusively in the germ cell fraction (i.e., PGCs), consolidating that recombined barcodes in gonads have derived specifically from germ cells in both sexes. (**F**) The total number of PGCs (or gonocytes/prospermatogonia), summed from the bilateral testes of *Prdm14-MiCM*;*Polylox*;*Dppa3*-*ECFP* mice, at E12.5 (with or without 4-OHT administration) and E13.5 (without 4-OHT administration). PGCs were counted based on immunostaining of Dppa3-ECFP and DDX4, and image analysis of encircled nuclei (see Methods). Horizontal bars indicate medians. No statistical significance was detected (two-sided MWW). (**G**) Three-dimensional representative images of E12.5 and E13.5 testes showing PGCs immunostained for Dppa3-ECFP and DDX4, visualized using ClearVolume^52^. Scale bar, 100 μm.

**Figure S2.**
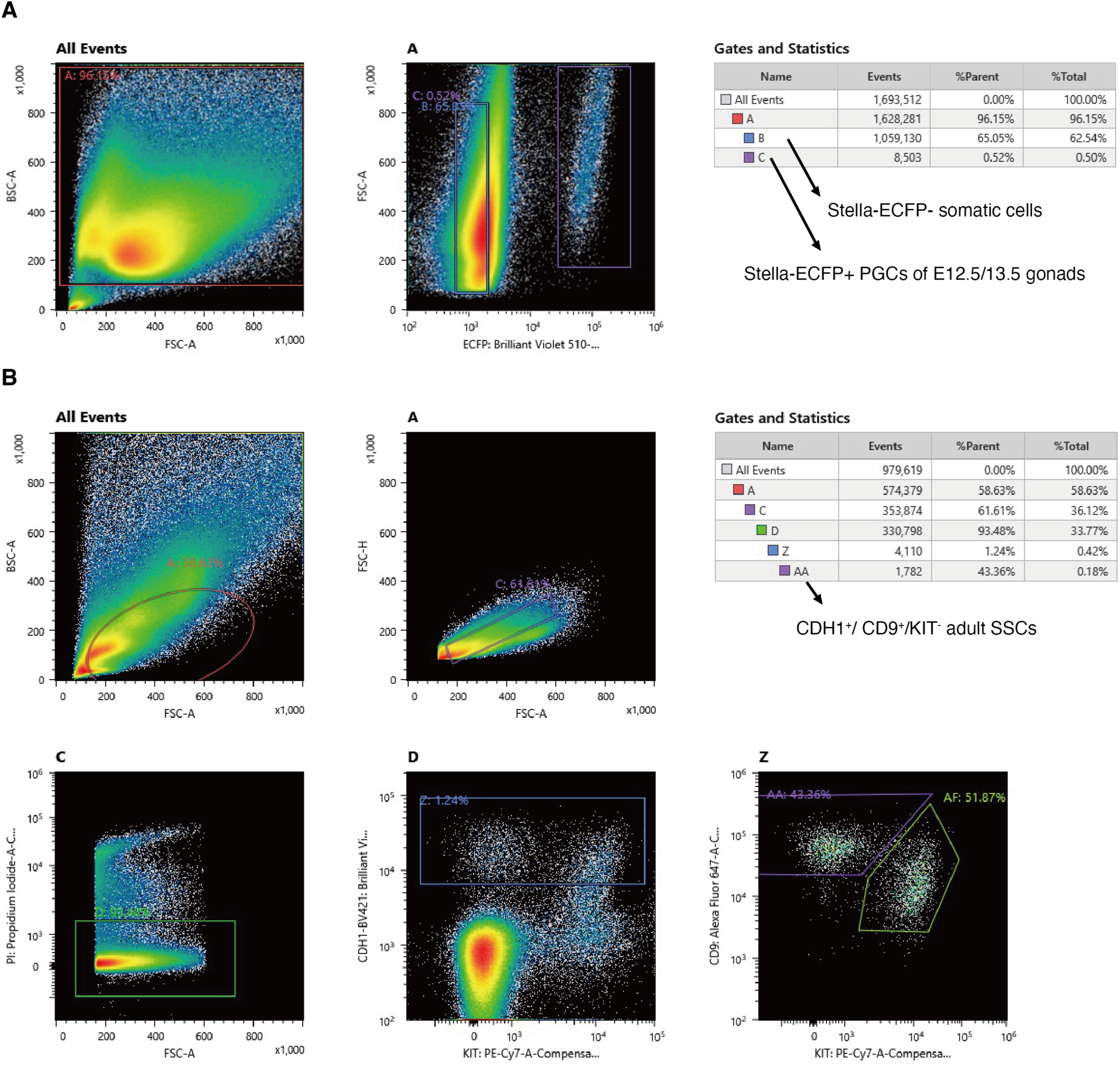
Gating strategy. (**A**) Gating strategy for embryonic gonadal PGCs and somatic cells. Cells with extreme FSC-A and BSC-A values were first excluded, after which the Dppa3-ECFP⁺ and ECFP⁻ fractions were independently sorted. (**B**) Gating strategy for adult SSCs. Cells were first gated for moderate FSC-A and BSC-A values, proportional FSC-A and FSC-H to select single cells, and low PI incorporation to exclude dead cells, followed by sorting for CDH1⁺ / CD9⁺ / KIT⁻ cells.

**Figure S3.**
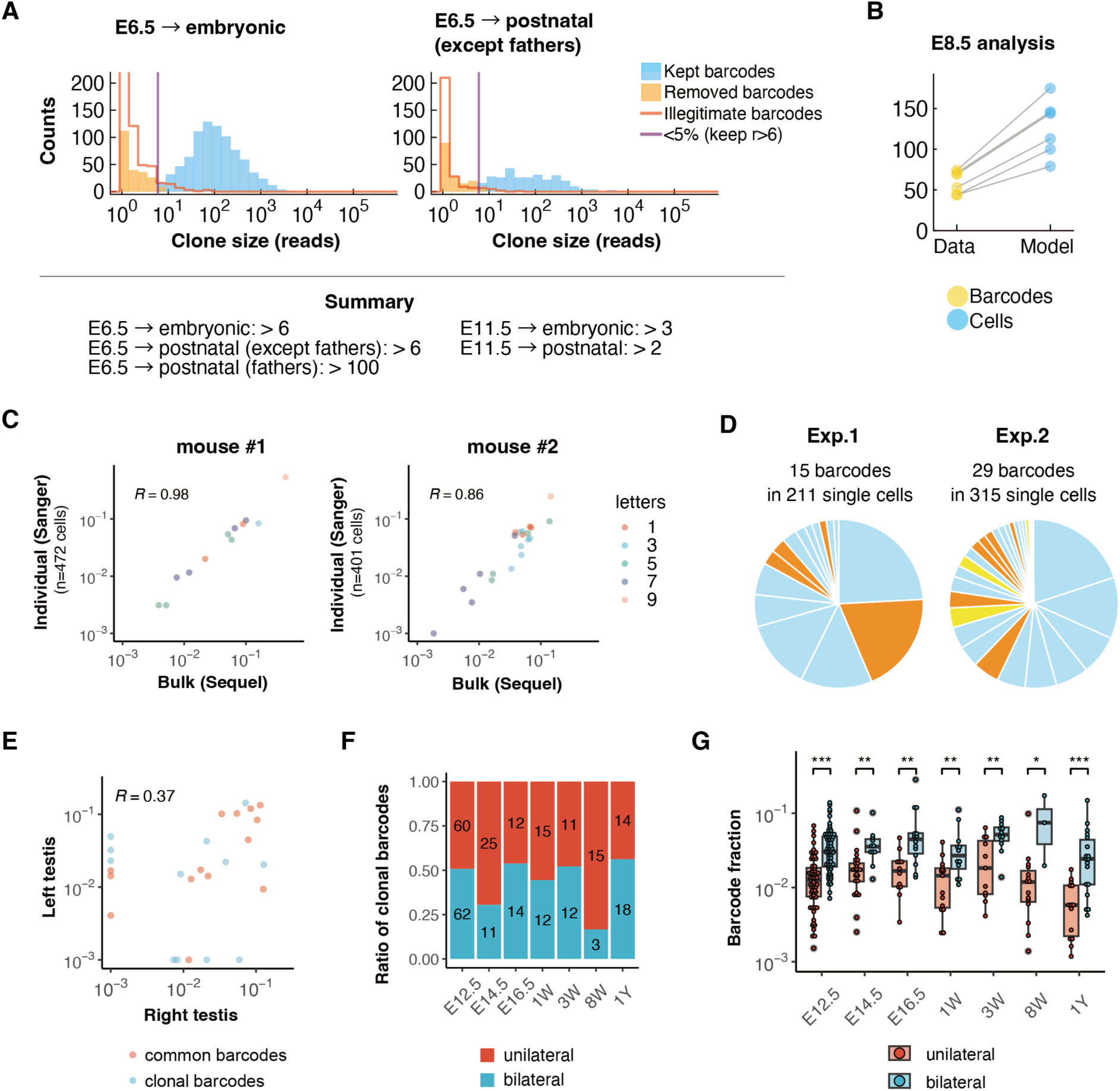
Barcode analysis of E6.5-labeled PGC clones. (**A**) Clone size (sequence read) distributions and criteria for barcode filtering. Barcodes were kept only if their clone size exceeded a defined threshold. The threshold (purple line) was set as the smallest clone size for which the number of remaining illegitimate barcodes (red) fell below 5% of the number of remaining legitimate barcodes (blue). Legitimate barcodes with clone sizes smaller than or equal to the threshold were excluded from subsequent analyses (orange). (**B**) Estimation of the cell number leading to the observed barcode outcomes, inferred from a model simulation with a given number of cells recombining their *Polylox* cassette intrinsically. The model was fitted to find the most probable cell number (blue) best explaining the unique barcodes observed in E8.5 PGCs, after induced at E6.5 (yellow). (**C**) Validation of proportionality between the bulk barcode reads by amplicon sequencing using long-read sequencers (Sequel, Pacific Biosciences) and the actual composition of barcoded cells. A CDH1^+^/CD9^+^/KIT^−^undifferentiated spermatogonia fraction (i.e., SSCs) were FACS-sorted from bilateral testes of two male mice, each barcoded at E6.5 and collected at 26 and 37 weeks of ages. On one hand, the *Polylox* sequences contained in this fraction were read by a bulk amplicon sequencing using Sequel; in parallel the same cell fraction was further sorted into individual cells, whose barcodes were amplified and read by Sanger sequencing. As displayed, the results of bulk sequencing showed strong linear correlation with individual read, irrespective of the letter length (color-indexed). 10^−3^ was added to each barcode’s frequency for log_10_ transformation; Spearman’s rank correlation coefficients (R) are shown. (**D**) Composition of barcodes induced at E6.5 and analyzed in E13.5 testes, through Sanger-sequencing the individually sorted PGCs. Results from two independent experiments were shown, with clonal barcodes indicated (*P_gen_*<0.001 and <0.0011, orange and yellow respectively). These results consolidate the conclusion based on the bulk sequencing that single PGC-derived clones induced at E6.5 unequally expand during their migratory stage. (**E**) Barcode composition included in right and left testes, with common (*P_gen_*≥0.001) and clonal (*P_gen_*<0.001) barcodes colored in red and cyan, respectively. Representative results from a single mouse (ID: m060) were shown. 10^−3^ was added to each barcode frequency for log_10_ transformation; Spearman’s rank correlation coefficient (R) is shown. (**F**) Fractions of clonal barcodes observed unilaterally versus bilaterally, collected from all mice harvested after E12.5. (**G**) Contributions of unilaterally versus bilaterally detected clonal barcodes in all recombined barcode reads. Note that bilaterally observed barcodes show significantly higher contributions at all time points. *** P < 0.001, ** P < 0.01, * P < 0.05, two-sided MWW.

**Figure S4.**
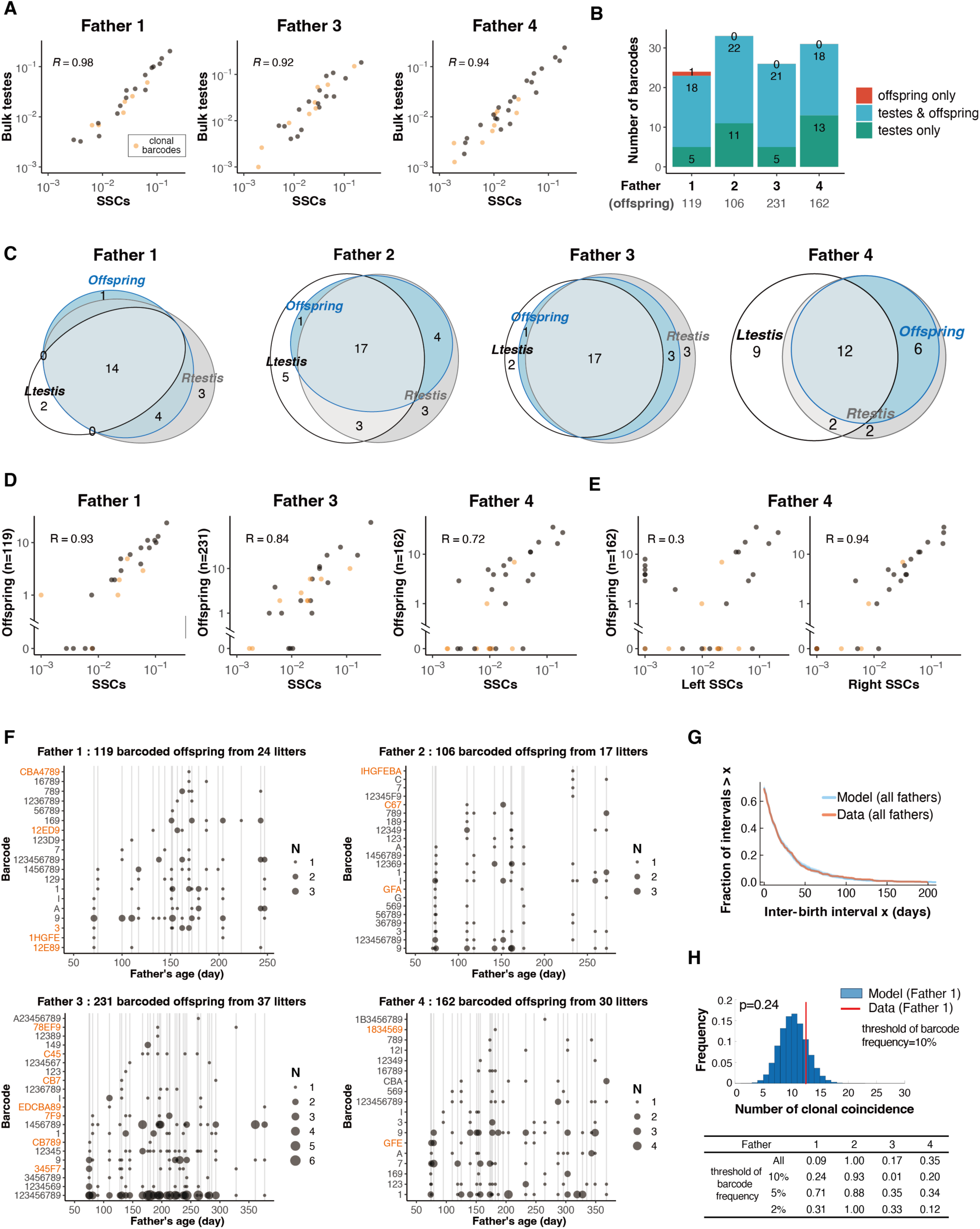
Barcode transmission from SSCs through spermatogenesis to the offspring. Supplementary data for Figure 2, involving four male mice barcoded at E6.5, mated with multiple females, and had their testes sampled at 1 year of age; followed by barcode reading in their SSCs, bulk testes, and offspring. (**A**) Correlation between the barcode compositions in the SSCs and that in the bulk testes for three fathers but Father 2 presented in Figure 2B. Note the similarly strong linear correlations. 10^−3^ was added to each barcode frequency for log_10_ transformation; Spearman’s rank correlation coefficient (R) is shown. (**B**) A bar plot showing diversity of barcodes appeared in fathers’ testes only (green), both father’s testes and offspring (cyan), and offspring only (red), summarized for each of the four fathers. All but 1 barcode identified in offspring were also found in the testes. The exceptional one likely derived from a PGC clone that became extinct before Father 1 reached one year of age. The number of offspring carrying barcodes are indicated below. (**C**) Venn diagrams indicating barcode overlapping among the left testis (white), right testis (gray) and offspring (cyan), summarized for each father. Note that all barcodes that appeared in Father 4’s offspring were included in Father 4’s right testis. (**D**) Correlation between barcode composition in SSCs and in offspring, with the result from Father 2 shown in Figure 2C. Note the similarly strong linear correlations. 10^−3^ was added to each barcode frequency for log_10_ transformation; Spearman’s rank correlation coefficient (R) is shown. (**E**) Correlation between barcodes found in SSCs from left or right testis of Father 4 and those observed in its offspring. Note that barcodes in offspring show strong correlation with SSCs from the right, but not the left, testis, suggesting that offspring of Father 4 originated exclusively from the right testis. (**F**) Summary of the barcodes transmitted to offspring, plotted for the father’s age on offspring’s birthday. Gray vertical lines mark each litter’s birthday; dots represent littermates whose size indicates the number of littermates sharing the same barcode. Clonal barcodes are colored in orange. (**G**) Analysis of inter-birth intervals in reference to a random-shuffle model. Inter-birth intervals were aggregated from each father, quantifying the time between births of pups with the same barcode. Interval times of zero arise from barcode coincidences in pups of the same litter. Model simulations were obtained by shuffling the barcodes of all pups for each father randomly, and aggregating intervals as done for the data. Solid lines represent data (red) and the median of repeated model simulations (blue); band indicates 95% credibility region of the model. (**H**) Probability of clonal coincidence, defined as offspring sharing the same barcodes in the same litter, were compared between model and data. The representative plot (top) shows the result using barcodes of Father 1 with frequency lower than 10%. Blue columns indicate the probabilities of the number of the clonal coincidence calculated by Monte Carlo samplings. Red lines indicate the actual number of clonal coincidences observed in the experiments. Probabilities based on different thresholds of barcode frequency are also shown (bottom table). These tests hardly showed statistical significance, consistent with the idea that each offspring was derived from PGC clones randomly selected from an SSC pool.

**Figure S5.**
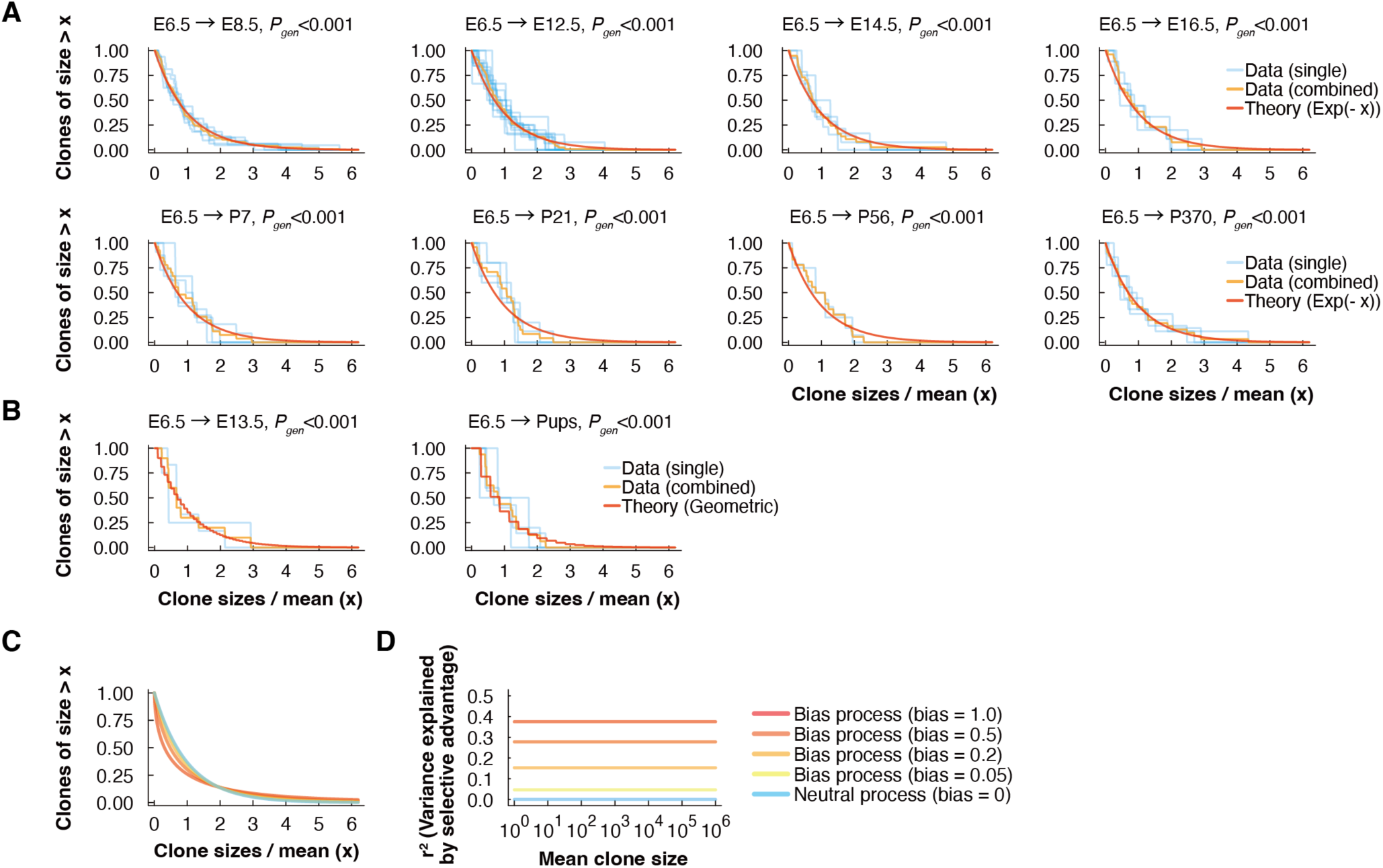
Rescaled clone size distributions conforming with neutral birth-death processes. (**A**) Size distributions of E6.5-barcoded PGC clones (i.e., read distribution of clonal barcodes with *P_gen_*< 0.001) at indicated time points were rescaled by the mean value for each mouse (blue) and grouped mice (orange), and were compared to the theoretical distribution expected from a neutral birth-death process following the continuous scaling law of the exponential function (red; Supplementary Text). (**B**) Size distribution of E6.5-induced PGC clones (with clonal barcodes) obtained from individual SSCs (analyzed at E13.5) and from individual pups, compared to the theoretical distribution predicted by a discrete geometric model assuming neutral clonal competition. (**C** and **D**) Effect of clonal bias and persistent selective advantages on clone size distributions. Rescaled clone sizes (C) are compared between the neutral model (blue), assuming neutral clonal competition without bias (birth-death models; A and B), with non-neutral dynamics at various bias levels (yellow to red). The bias parameter *b* is defined as CV_bias_ = (1 + *b*) CV_neutral_, i.e. enhancing the coefficient of variation (CV) of the neutral clone sizes. Levels of clonal bias correspond to a certain fraction of the clone size variability that is explained by selective advantages between clones, as quantified by the squared correlation *r*^2^ (D; Supplementary Text). It holds that *r*^2^ = A*b*(*b* + 2))/M2A1 + (*b*(*b* + 2))O, for any mean clone size. The remaining fraction of the clone size variability, 1 − *r*^2^, originates from neutral stochasticity. Clone size variability for *b* = 0.2 already deviates noticeably from the neutral case (C) and the data (A and B), indicating that PGC clone size variability is, by and large, driven by neutral stochastic effects (i.e., 85% – 100%).

**Figure S6.**
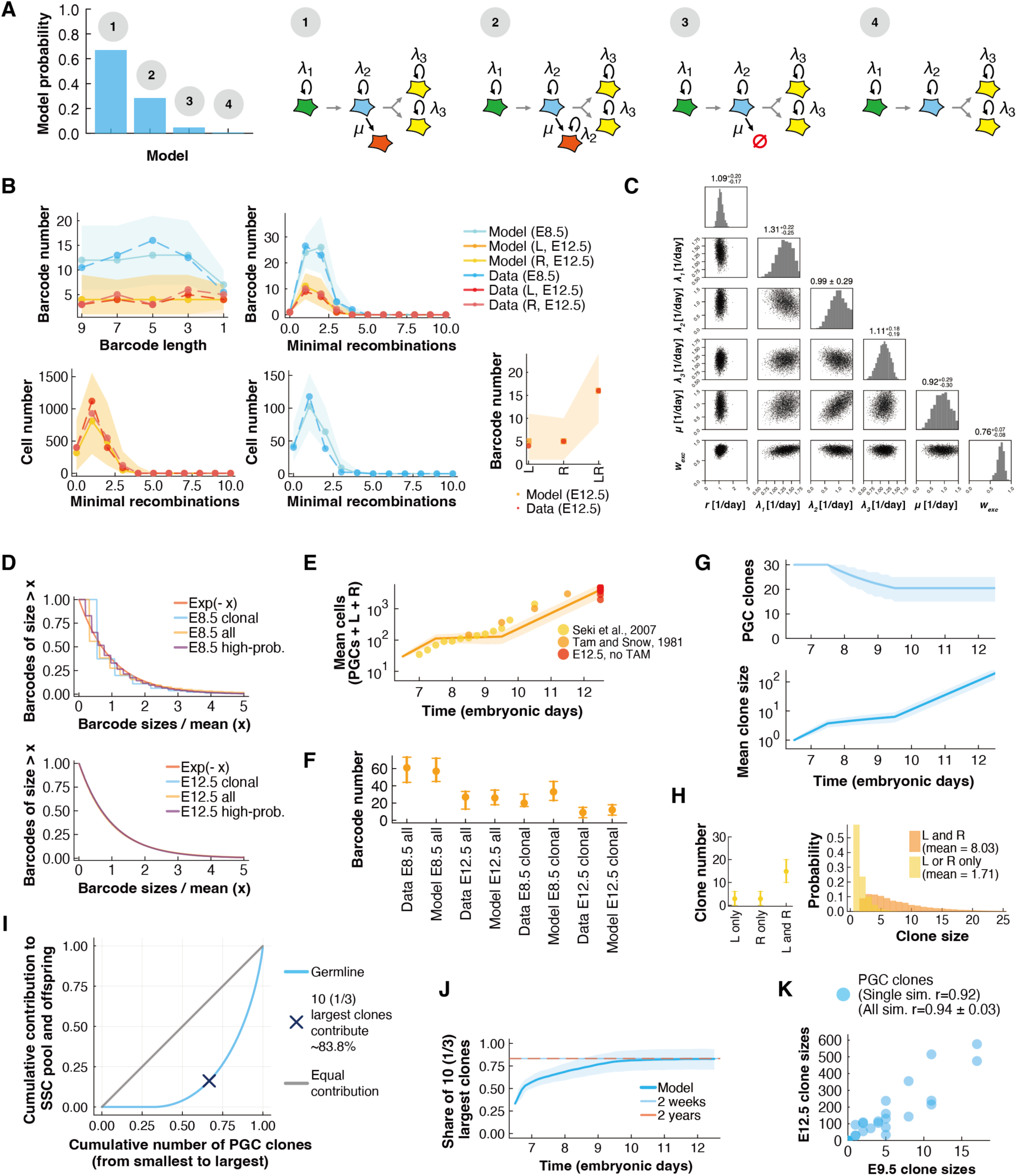
Modeling the clonal dynamics of PGCs from E6.5 to E12.5. (**A**) Posterior probabilities (left) inferred for the four tested modeling schemes (1-4; right). All models simulate PGC clonal dynamics in three sequential phases of birth-death processes from E6.5 to E7.5 (green), E7.5 to E9.5 (blue) and, after splitting into left and right gonads, from E9.5 to E12.5 (yellow), following the respective division rates (*λ_1_*, *λ_2_* and *λ_3_*). The top model (1) simulates PGC loss with a rate *μ* from the second-phase PGCs (blue) into a non-arriving PGC fraction (red) that is still measured at E8.5 but does not proliferate and does not reach the gonads (yellow). Model 2 is like model 1, except that the non-arriving PGC fraction continues to divide with the same rate. Model 3 simulates PGC loss as immediate cell death. The model (4) which assumed all PGCs to survive without loss showed the lowest predictive capacity. (**B**) Barcode and clonal dynamics statistics used for the inference. Posterior statistics of the top model (Model 1 in (A)) are shown with 95% credibility bands, compared to the data as indicated. (**C**) Posterior parameter distributions as inferred from the data for the top model. The set contains biological parameters specifying PGC division and loss (*λ_1_*, *λ_2_*, *λ_3_* and *μ*), and technical parameters specifying the *Polylox* recombination kinetics (recombination rate *r* and excision bias *w*_exc_). (**D**) Predicted distribution of *Polylox* barcode contribution simulated for E8.5 and E12.5 (top model with posterior parameters), rescaled with their means, based on clonal (blue), non-clonal/high-probability (purple) and all (orange) barcodes. In comparison, the theoretical scaling law of birth-death process (red) was indicated. The discreteness in E8.5 barcode sizes (upper panel) comes from the low cell numbers at this time point. Note the good agreement with the data, as shown in Figure S5 for clonal barcodes. (**E**) Predicted germ cell numbers (excluding non-arriving PGCs), shown as mean (lines) with 95% credibility bands. As reference, experimental measurements of embryonic PGC numbers are indicated (dot; see also Figure S1F). (**F**) Prediction of the total and clonal *Polylox* barcode numbers at E8.5 and E12.5 inferred by the top model, compared with the observed data. Median number (dot) with 5^th^ and 95^th^ percentiles (error bars) are shown. Note that the number of all barcodes is fitted in (B), but the clonal barcode numbers are a model prediction. (**G**) Prediction of the clone dynamics originated from E6.5 PGCs based on the top model with inferred parameters. Number of surviving E6.5 PGC-derived clones (top) and mean clone sizes of surviving E6.5 PGC-derived clones (bottom). Both model predictions (lines) are shown with 95% credibility bands. (**H**) Predicted number and size distribution of clones originating from the initial ∼30 PGCs at E6.5, which arrived at the left (L) and/or right (R) gonads at E9.5, based on the top model with inferred parameters. Median clone number (dot) with 2.5^th^ and 97.5^th^ percentiles (error bars). The model predicts that left-right shared clones are bigger E9.5 and in the long terms, consistent with the experimental data (see also Figures 3B and S3G). (**I**) Skewness in the male germline (blue) as quantified by the combined number of E6.5 PGCs (*x*-axis) reaching a certain share in the germ cell pool from around ∼E11 onwards, including later share in SSCs, sperm and offspring (*y*-axis); akin to Lorenz curves for economic wealth or income distributions. The Lorenz curve incorporates effects of non-arriving PGCs (∼32%) that do not contribute at all and uneven clone sizes among the survivors as set by neutral drift (exponentially distributed). A hypothetical equal contribution of PGCs is indicated (gray). Specific case of the ∼10 largest clones indicated (dark blue marker), see also (J). (**J**) The share of the ∼10 (33.3% of initial ∼30) largest clones among the initial E6.5 PGCs over time, developing from a combined effect of PGC loss and clone size divergence. The share stabilizes from ∼E11 onwards (Figure 3C), reaching levels of ∼84%, as seen 2 weeks and 2 years after birth (as indicated; using extended lifetime model, Figure 5) and eventually in offspring due to proportional clone-size dependent transmission (Figure 2); see also (I). (**K**) Correlation between the predicted clone sizes of the same clones between E9.5 and E12.5; a representative simulation (sim.) shown, correlation values indicated for the single and all simulations.

**Figure S7.**
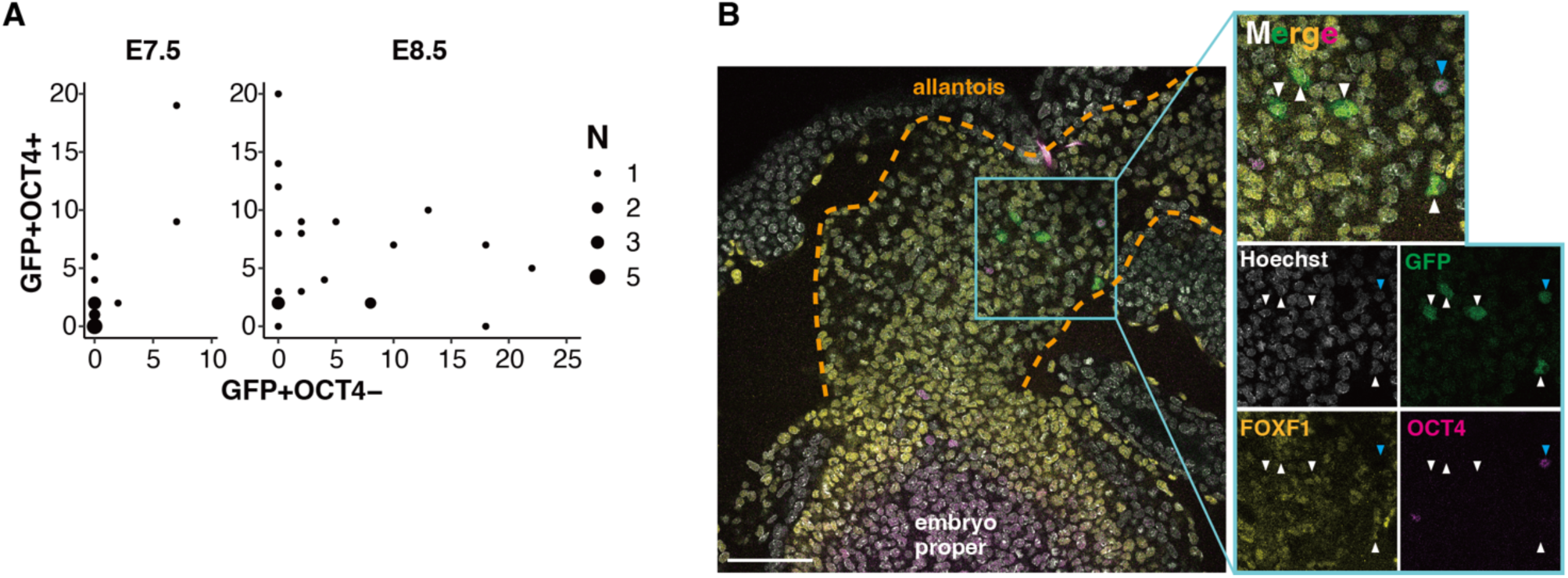
Distribution of E6.5-labeled GFP^+^ PGC descendants. (**A**) The numbers of GFP^+^OCT4^+^ and GFP^+^OCT4^−^ cells in E7.5 (left) and E8.5 (right) embryos. Dot size indicates the number of embryos with the same cell counts. (**B**) Representative images of E8.5 embryo stained for GFP, OCT4, and FOXF. Cyan and white arrowheads indicating GFP^+^OCT4^+^ and GFP^+^OCT4^−^ cells, respectively. All four GFP^+^OCT4^−^ cells were positive for FOXF1, a marker of extraembryonic mesoderm including allantois^19^. Scale bar, 100μm.

**Figure S8.**
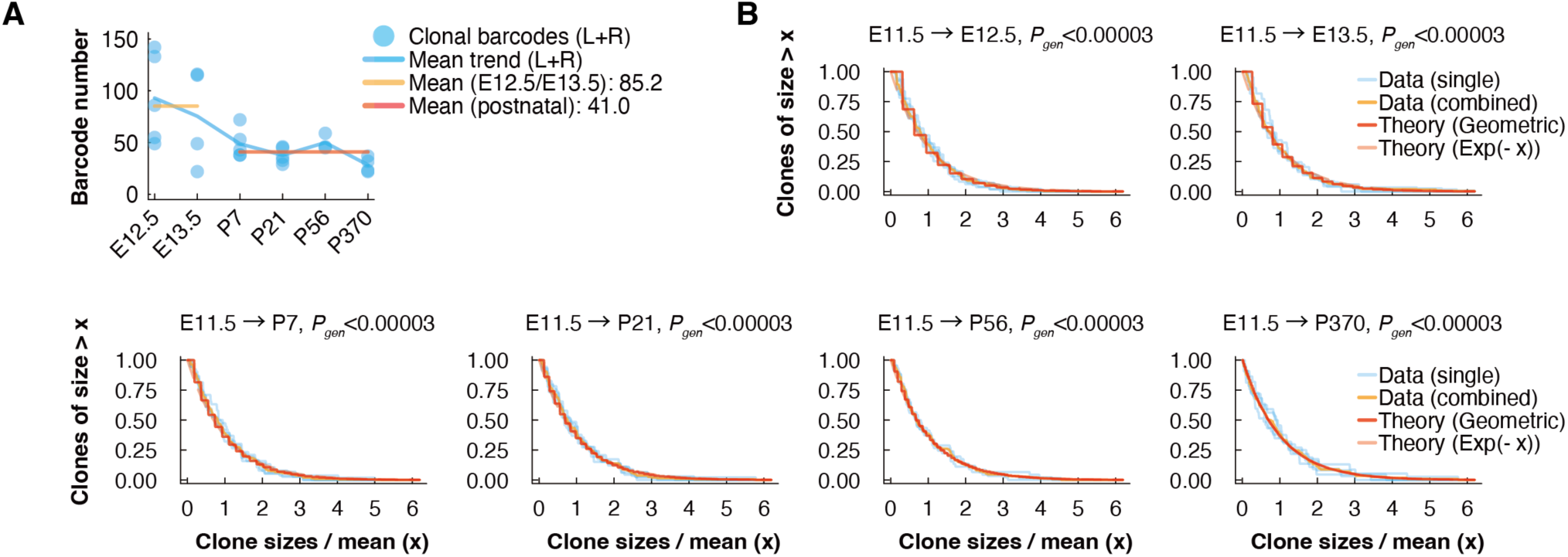
Estimating the survival chance of single E11.5/E12.5 PGC-derived clones. (**A**) Number of clonal barcodes (*P_gen_* < 3 × 10^−5^; left and right testes combined) induced at E11.5 and recovered over embryonic and postnatal stages as indicated. The reduction in the mean number of clonal barcodes (horizontal lines) from ∼85 in E12.5–13.5 to ∼41 after birth indicates that the survival chance of a single PGC-derived clone from E12.5 to adult stages is approximately 50%. See Figure 4B for total barcode numbers. (**B**) Size distributions of E11.5-barcoded PGCs (*P_gen_* < 3 × 10^−5^; left and right testes combined) at indicated time points were rescaled by the mean value for each mouse (blue) and grouped mice (orange), and were compared to the theoretical distributions expected from a neutral birth-death process (Supplementary Text), in form of the continuous scaling limit of the exponential function (light red, dashed line) and the geometric distribution accounting for the discreteness of small read numbers (red, solid line).

**Figure S9.**
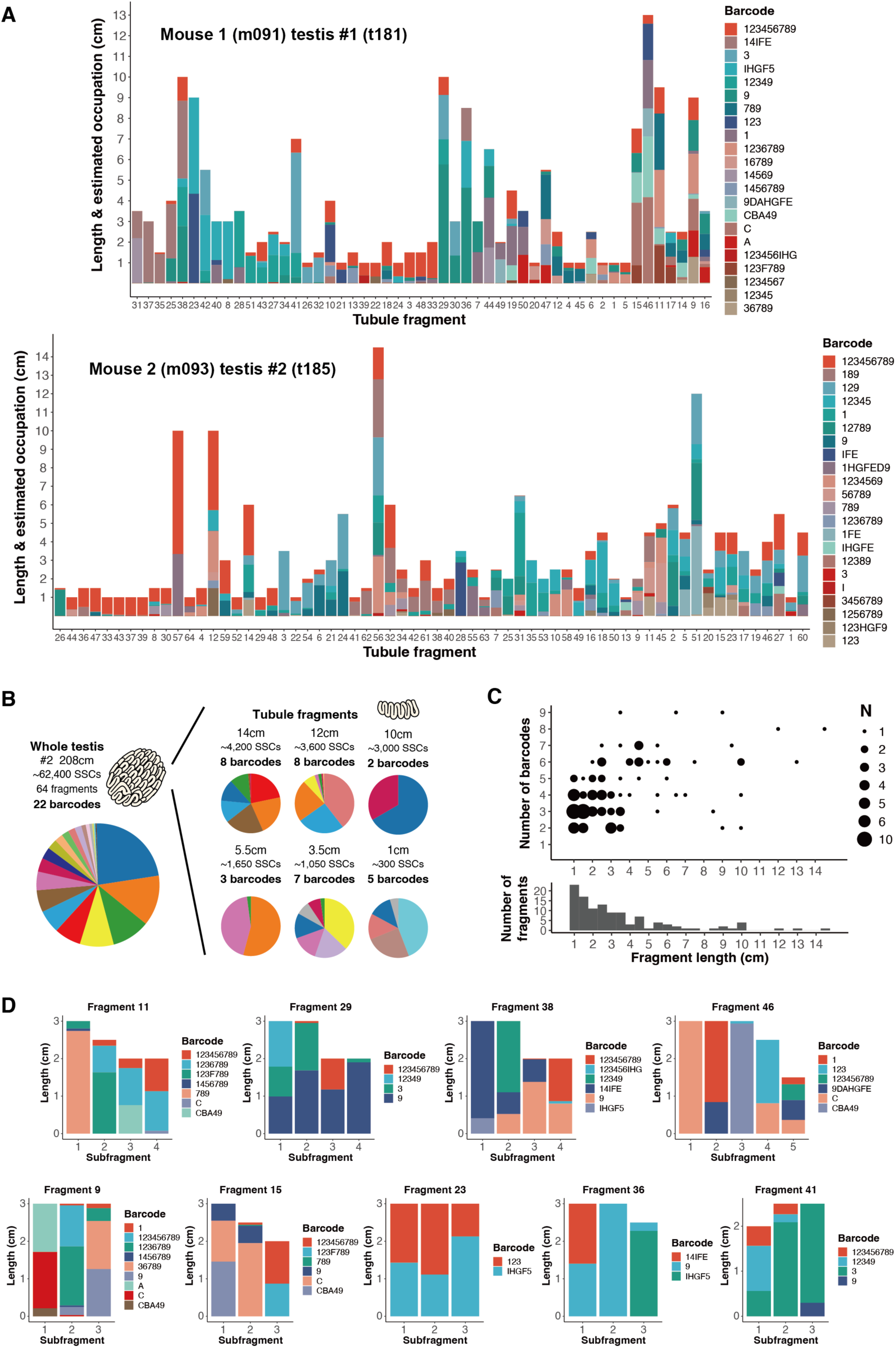
Barcode distribution over the seminiferous tubules in the adult testis. (**A**) Summary of two sets of barcode reads, each obtained from seminiferous tubule fragments representing a whole single testis each, separately sampled from two different individuals that were barcoded at E6.5 and sampled in 8 (Mouse 1) and 9 (Mouse 2) weeks of age. Each column represents the set of barcodes observed in a single fragment of the indicated length, color-coded according to the identity and composition of barcodes, as specified on the right. Fragments are aligned on the x-axis according to hierarchical clustering of barcode frequency. (**B**) Summary of the uneven barcode distribution of the testis #2 from Mouse 2, presented in a same manner with Figure 4, D and E. (**C**) Number of barcodes detected in each fragment with different lengths (shown at the bottom). Data from the two testes (#1 and #2) are combined. Dot size indicates the number of fragments with the same length and barcode count. (**D**) Barcode compositions in long contigs of tubule fragments obtained from testis #1 of Mouse 1, analyzed separately in their sub-fragments. Sub-fragments are numbered in spatial sequence, except for the branched fragment (#46). Each column depicts each sub-fragment, with the total heights reflecting the tubule lengths and colors indicating the barcode identities (specified for each panel).

**Figure S10.**
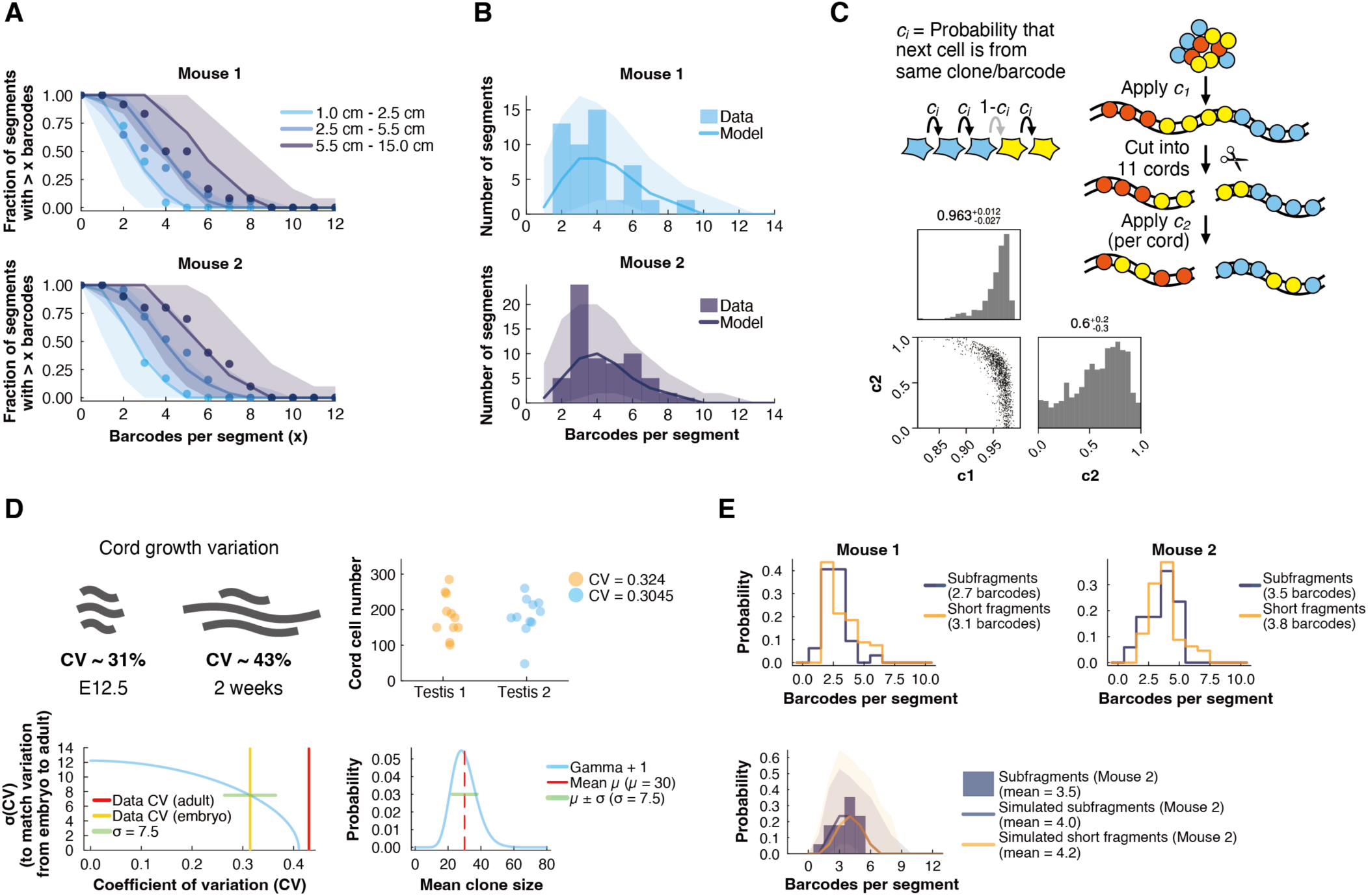
Inference of a model for tubular PGC dynamics. Setup and inference of a model extending the PGC clone dynamics after E9.5, including phases of testis cord formation at ∼E12.5, their expansion (E12.5 to 2 weeks), followed by homeostasis as analyzed experimentally at 8 weeks. (**A**) Statistics used for the model inference. Distributions of *Polylox* barcode number included in tubule segments (Figure S9) were fitted simultaneously for three categories based on the indicated segment lengths for each of the two testes from two mice analyzed (barcoded at E6.5, simulated until 8 weeks after birth). Data (dots) and posterior model inference (solid lines with 95% credibility bands) shown. (**B**) Predictive capacity of the model validated in term of the distribution of barcode numbers across all analyzed segments (data not directly used for fitting). Data (histograms) and the model prediction (solid line with 95% credibility bands) are overlayed. (**C**) Model setting of distributing the PGC clones upon compartmentalization through testis cord formation at E12.5. The parameters *c_1_* and *c_2_* model the spatial distribution of clones on the forming ∼11 tubule loops (effectively one-dimensional) at E12.5. Both parameters measure how strongly cells of the same clone form local clusters (top schemes). The first parameter *c_1_* controls the cluster sizes of clones within a cord at the time of compartmentalization, and with that, the number of clones per cord. When PGCs subsequently align into the one-dimensional tubular arrangement, clusters may mix and yield the final organization of local patches. The sizes of these local patches are controlled by the second parameter *c_2_* acting on the cluster sizes. Inferred posterior model parameters *c_1_* and *c_2_* (lower left). (**D**) Prediction of the variation in the extension of testis cords/seminiferous tubules within a single testis between E12.5 and 2W after birth. Summary of the coefficient of variation (CV) of cord cell numbers at E12.5 and in 2W after birth (top left), based on our measurement (see the top right panel) and Nakata et al., 2015^32^, respectively. Numbers of germ cells included in individual testis cords of two E13.5 testes, with respective CVs indicated (top right). Shown values were rescaled to match the total germ cell number at E12.5 (∼2,000) using E13.5 data (Figures S1F and S1G) as a close approximation. Inferring the cord variation parameter σ (i.e., the variability of extension of individual testis cords/seminiferous tubules within a single testis between E12.5 to 2W; bottom left). The blue line indicates the required heterogeneity to increase the initial embryonic CV to the adult CV during expansion. Best prediction of the 2W value from of the E12.5 value is given with σ ≈ 7.5. (see Supplementary Text). A gamma distribution was employed to simulate cord growth with the inferred cord variation parameter of σ ≈ 7.5 (Bottom right). (**E**) Validation of our modeling scheme of heterogeneous extension of testis cords/seminiferous tubules (with σ ≈ 7.5) by independent, barcodes-based data. Heterogeneous cord expansion (as specified by CVs in (D)) predicts a shift in the mean barcode number between short fragments and sub-fragments (pieces of a long fragment) when accounting for an identical underlying fragment length distribution. Data (top panels; means indicated in legend) show a slight shift of ∼0.3 barcodes, consistent with the model prediction (lower panel) by ∼0.2 barcodes (see Supplementary Text for more details).

**Figure S11.**
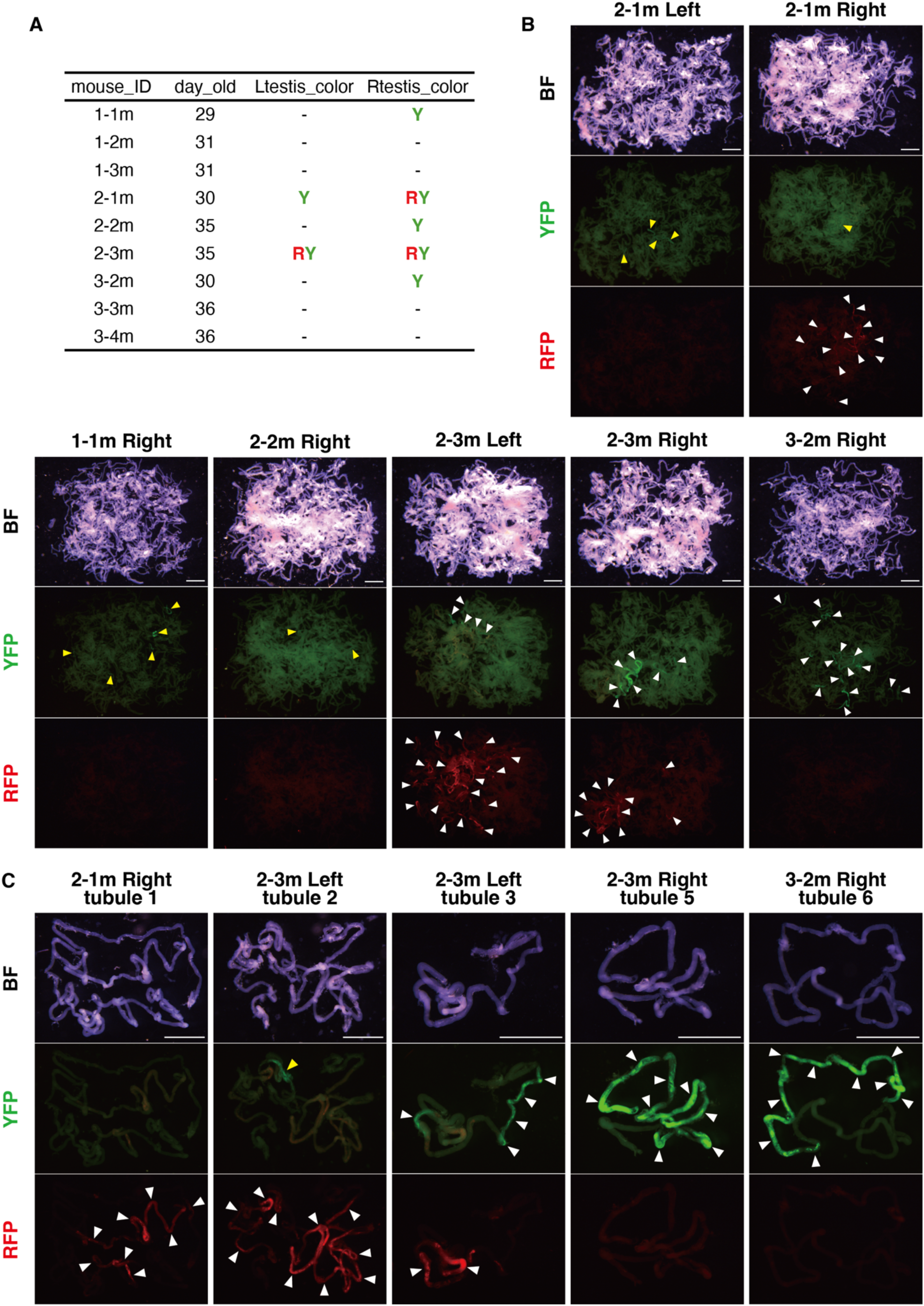
Visualization of E6.5 PGC-derived clones in adult using *Confetti* reporter, related to Figure 5C. Following the experimental design presented in Figure 5C top panel, PGCs were sparsely labeled at E6.5 and their clones were visualized at 4-5W after birth. (**A**) Summary of the analyzed nine *Prdm14-MiCM*;*Confetti* mice and their fluorescent cells in their testes. R and Y indicate cRFP and cYFP, respectively. Note that only 5 in 9 mice contained fluorescent cells, and 4 of them having colors found only in unilateral testis, suggesting highly oligo-clonal labeling. mCFP and nGFP patches were not observed. (**B**) Thoroughly disentangled whole seminiferous tubules in testes of *Prdm14-MiCM*;*Confetti* mice. Seven testes in (A), which included one or more fluorescent-labeled patches are shown. Note the prominent green patches in 2-3m Left, 2-3m Right, and 3-2m Right, and red patches in 2-1m Right, 2-3m Left, and 2-3m Right testes (white arrowheads). Small green patches are also found, obscured by background signals in 1-1m, 2-1m and 2-2m (yellow arrowheads). (**C**) Higher magnification views of isolated fragments of seminiferous tubules carrying patches of fluorescence-labeled cells prepared from the indicated testes. Note prominent and extended green and red patches in tubules 3, 5, 6 and tubules 1, 2, 3, respectively (white arrowheads), and small green patches in tubules 2 (yellow arrowhead). Scale bar, 2mm.

**Figure S12.**
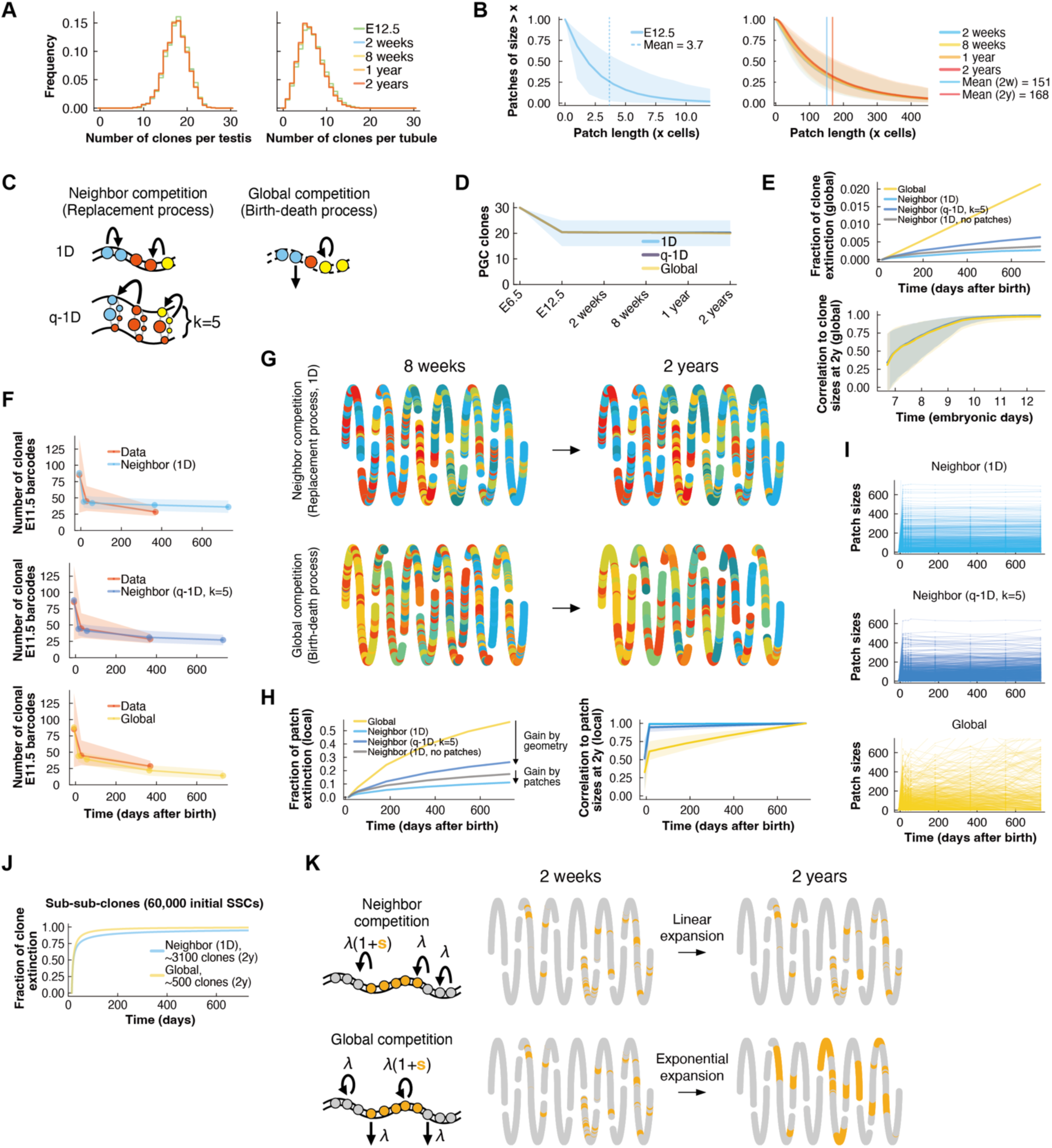
Inferred spatiotemporal dynamics of PGC clones over the male life span. Lifelong dynamics of male germline clones (instead of *Polylox* barcodes) from their emergence at E6.5 to 2 years of age were inferred both globally (averaged over an entire testis) and locally (considering spatial organization within seminiferous tubules). The integrative model consists of a sequence encompassing early clonal dynamics from E6.5 to E12.5, the formation and elongation of testis cords/seminiferous tubules by 2 weeks after birth, and an extended homeostatic phase lasting up to 2 years of age. (**A**) Model prediction for the number of surviving PGC clones within one testis (left) and within one of the ∼11 testis cords/seminiferous tubules (right), at different time points. (**B**) Model prediction of patch size distributions at E12.5 (left) and at multiple postnatal time points (right). Median prediction (solid lines) and 95% credibility band are shown, with mean patch sizes indicated by vertical lines. Homeostatic phase was modeled under neighbor competition dynamic within one-dimensional tubules. (**C**) Schemes of regulation modes during the adult homeostatic phase. Neighbor competition in tubules is compared to a hypothetical global competition model as reference, to investigate the effect of the tubular geometry on clone and patch size dynamics. In the scheme of neighbor competition (left), a lost stem cell through differentiation is replaced by one of the neighbors to maintain a constant stem cell pool. A one-dimensional (1D) geometry (left top), in which a lost stem cell is replenished by duplication of either the left or right neighbor. A quasi-one-dimensional (q-1D) tubular configuration (left bottom) with k=5 cells in the circumference (based on Klein et al., 2010^31^), in which a lost stem cells is replenished by duplication of any one of the 4 neighbors (left, right, bottom or top). In the hypothetical global competition mode (right; modeled as a critical birth-death process), stem cell loss and duplication occur stochastically and without spatial coupling, while maintaining a balanced total number of SSCs. (**D**) Model prediction for the number of surviving E6.5 PGCs (starting from ∼30) up to 2 years (solid lines; 95% credibility band for 1D). Note that after E12.5, global PGC clones become very stable, in line with the observed *Polylox* barcodes induced at E6.5 (Figure 1). Clone stability across the testis is robust, independent of the mode of adult homeostatic regulation (shown in (C)). Detailed dynamics for the early phase (E6.5 to E12.5) are shown in Figure S6G. (**E**) Model prediction for extinction probabilities and size correlation of global PGC clones following different modes of homeostatic regulation shown in (C). Over the testis wide, PGC clones show high stability from as early as ∼E11, as indexed by low extinction during postnatal phase (top) and high correlation to their long-term clone sizes at 2 years (bottom), with the neighbor competition modes showing higher stability. Model bands show 95% credibility regions. (**F**) Projection of the model dynamics by following the E12.5 germ cell-derived clones (PGC sub-clones) for their survival over time (compared for the three homeostatic regulation modes) with those observed in the E11.5 barcoding data (clonal barcodes, *P_gen_* < 3 × 10^−5^). The q-1D setup in the neighboring competition yielded the highest predictive power, likely due to the markedly smaller size of E12.5 germ cell-derived clones compared to those from E6.5 PGCs. Data and model bands show 95% credibility. (**G**) Model prediction of the spatial organization of PGC clones in postnatal seminiferous tubules of a single testis from 8 weeks to 2 years of age, under different homeostatic regulation either by neighbor competition within tubules (upper panels) or by a hypothetical global competition as a reference (lower panels). In both scenarios, the testis is modeled as comprising 11 separate tubule loops, which contain a one-dimensional array of ∼60,000 SSCs in total color-coded by PGC clone identity (one of the ∼30 initial clones). Cells belonging to the same clone (indicated by the same color) often appear repeatedly as separate patches, within and across the tubule loops. Note the spatial organization of such patches is high stable under neighbor competition, but unstable under a global competition. (**H**) Model prediction for extinction probabilities and size correlation of local patches, following different modes of homeostatic regulation shown in (C). The high stability of the neighbor competition dynamics is more clearly underscored by the fates of local patches, as evidenced by the extinction rates (left) and correlations with long-term patch sizes (right). Model bands show 95% credibility regions. (**I**) Model simulations of individual patch sizes over a testis, comparing different modes of homeostatic regulation. Note that global competition (birth-death process) shows lower stabilities as well. (**J**) Model prediction for the fraction of extinction of single SSC-derived clones (PGC sub-sub-clones) during homeostasis from 2 weeks to 2 years postnatally, comparing neighbor competition in a 1D tubular geometry with global competition. Neighbor competition maintains a higher clonal diversity after 2 years (indicated in legend), starting with ∼60,000 SSC clones at 2 weeks. (K) Model prediction for the expansion of a single E6.5 PGC clone with a selective advantage during postnatal homeostasis, comparing neighbor competition in a 1D tubular geometry (top) with global competition (bottom). An identical clonal arrangement at 2 weeks (left) was simulated up until 2 years (right). The selected clone (orange patches) was equipped with an advantage of *s* = 1%, increasing the rate of cell division (λ) in contrast to neutral PGC clones (all shown in gray). Under neighbor competition, enabled by the tubular geometry, the selected clone was effectively confined; in contrast, the clone expanded exponentially under the hypothetical global competition (Supplementary Text).

**Table S1. (separate file**)

List of mice subjected to barcode bulk sequencing in this study.

**Table S2. (separate file)**

Number of testicular PGCs and PGCs within E13.5 testis cords, quantified by image analysis.

**Table S3. (separate file)**

Barcodes detected in E8.5 embryos and in FACS-sorted adult bulk SSCs after E6.5-barcoding.

**Table S4. (separate file)**

Barcodes detected in bilateral testes after E6.5-barcoding.

**Table S5. (separate file)**

Barcodes detected in individual E13.5 PGCs or adult SSCs.

**Table S6. (separate file)**

Barcodes detected in individual offspring of E6.5-barcoded males.

**Table S7. (separate file)**

Number of GFP+ cells in E7.5 and E8.5 embryos.

**Table S8. (separate file)**

Barcodes detected in bilateral testes after E11.5-barcoding.

**Table S9. (separate file)**

List of seminiferous tubule fragments analyzed in E6.5-barcoded mice.

**Table S10. (separate file)**

Barcodes detected in seminiferous tubule fragments

